# Not all West Nile virus lineages behave alike: vector competence and minimum infectious dose differences of lineages 1 and 2

**DOI:** 10.64898/2026.02.03.700853

**Authors:** Patrick Höller, Felix Gregor Sauer, Renke Lühken, Norbert Becker, Jonas Schmidt-Chanasit, Stephanie Jansen, Anna Heitmann

**Author notes:** Corresponding author (Bernhard-Nocht-Strasse 74, 20359 Hamburg, Germany;).

## Abstract

The globally distributed arbovirus West Nile virus (WNV) continues to expand across Europe, with rising numbers of human cases and an increasingly broad geographic distribution. WNV is primarily transmitted by mosquitoes of the genus *Culex*. Out of nine WNV lineages, human pathogenicity has been clearly established for lineages 1 and 2, but differences in their transmission dynamics, such as minimal infectious dose and transmission efficiency, remain poorly understood. In this study, we investigated how viral lineage, mosquito species, and blood meal titer influence the vector competence of two primary WNV vectors from Europe, *Cx. pipiens* biotype *pipiens* and *Cx. torrentium*, as well as the invasive mosquito species *Aedes albopictus*. Female mosquitoes were fed with increasing blood meal titers containing either of the WNV lineages. After an incubation period of 14 days at a mean temperature of 24°C, mosquito body titers were quantified, and the presence of infectious viral particles in the saliva was assessed. Our results revealed clear differences between the two lineages. Lineage 2 resulted in higher transmission efficiencies across all three species and required lower infectious doses to cause transmission. Among the tested species, *Cx. torrentium* proved to be a highly competent vector (max. transmission efficiency = 30%, minimum infectious dose = 10^5^ TCID_50_/mL), despite its underrepresentation in research. These findings provide detailed insights into how viral lineage, mosquito species, and blood meal titer might shape WNV transmission, informing future risk assessments and efforts to mitigate WNV transmission in Europe.

## Introduction

West Nile virus (WNV), species *Orthoflavivirus nilense*, within the family *Flaviviridae* and the genus *Orthoflavivirus*, is an emerging mosquito-borne virus of public health concern in Europe [1,2]. Over the last decade, several outbreaks have been recorded, and the affected area has expanded into Central Europe, including Germany. The spread of WNV in Europe can, in large parts, be attributed to higher temperatures due to the ongoing global warming, which, among other effects, shortens the viral extrinsic incubation period. The geographic range of the virus is expected to expand further and could place up to 244 million people across Europe at risk of infection in the future [2,3]. Approximately 20% of WNV infections are symptomatic, causing West Nile fever, characterized by headache, fever, myalgia, and fatigue. Less than 1% develop neuroinvasive disease, such as meningitis or encephalitis, with a case fatality rate of up to 17% [4]. Vaccines for horses have been available since the early 2000s, but no human vaccine has progressed past phase II [5]. The largest epidemic recorded in Europe occurred in 2018, with 1548 confirmed autochthonous human cases and 166 deaths across eleven countries [6]. The most recent data from 2025 reports once again above-average numbers with 989 cases and 63 deaths in 14 countries [7].

WNV is maintained in an enzootic cycle between mosquitoes as vectors and reservoirs, and birds as amplifying hosts. Mosquitoes of the genus *Culex* are considered the main vectors of WNV, with *Cx. pipiens* s.s. being the most abundant in Europe. *Culex pipiens* s.s. can be further subdivided into two distinct biotypes: *pipiens* and *molestus* [4,8,9]. Recent data show that both biotypes, as well as *Cx. torrentium*, feed on avian, human, and nonhuman mammalian hosts [10]. In Central and Northern Europe, *Cx. torrentium* is predominant, but often not distinguished from *Cx. pipiens* s.s., despite its epidemiological importance [11]. Another species of interest is *Ae. albopictus*, the most widespread invasive mosquito in Europe. Although not considered an important WNV vector, the species is competent for WNV and exhibits an opportunistic feeding behavior, warranting further research into its role in the WNV transmission cycle [12].

WNV can be subdivided into nine distinct lineages, six of which have been detected in Europe. WNV lineage 1 (L1) and lineage 2 (L2) are most frequently associated with diagnosed infections in humans and animals [13]. WNV-L1 was first detected in Europe in the 1960s, presumably introduced by migratory birds, and has caused multiple outbreaks since. In contrast, WNV-L2 was first described in Europe in 2004, likely also introduced via migratory birds, and subsequently spread rapidly through Southern and Central Europe, where it has become the dominant cause of human cases [14]. However, WNV-L1 is still co-circulating in Europe, e.g. in Italy, and is the prevailing lineage in other parts of the world [13].

The virus can also be transmitted to mammalian hosts, including humans and horses [4]. However, humans and horses are considered incidental dead-end hosts, as their viremia is typically too low to further infect mosquitoes [4]. In contrast, viremia levels in avian species vary by several orders of magnitude and can reach levels of more than 10^8^ plaque-forming units (PFU) per mL, which influences their potential to be infectious for mosquitoes [15,16]. Therefore, mosquitoes encounter vastly different levels of viremia when feeding on animals, making it important to understand how viremia influences mosquito infection, and which viremia levels are sufficient to enable transmission (i.e. the minimal infectious dose required for transmission, MID). A lower MID implies that mosquitoes can become infectious after feeding from a broader range of hosts, as even closely related birds can have widely different peak viremias [15]. Interestingly, a recent report of a human co-infection with WNV lineages 1 and 3 described an unusually high WNV-L1 RNA concentration of 10^5.5^ PFU equivalents/mL, raising the question of whether humans can serve as amplifying hosts under very specific conditions [17].

In this study, we aimed to improve the understanding of the differences between WNV-L1 and L2 in terms of their infection dynamics in mosquitoes. We infected *Cx. pipiens* biotype *pipiens, Cx. torrentium*, and *Ae. albopictus* with WNV-L1 or L2 at varying blood meal titers. Using forced salivation assays, we assessed how blood meal titer, mosquito body titer, and mosquito species influence the infection and transmission.

## Methods

### Origins of mosquitoes, viruses, and cells

Three mosquito species were tested: *Ae. albopictus, Cx. pipiens* biotype *pipiens* and *Cx. torrentium*. The *Ae. albopictus* colony was established in 2016/17 from eggs collected in Freiburg, Germany. For both *Culex* species, egg rafts were field-collected in northern Germany (Lat: 53.467821, Long: 9.831346) in 2024 and 2025. Each egg raft was reared individually at 21°C, and five to ten larvae were used to molecularly determine the species and biotype, as described by Rudolf et al. [18]. Ten adults of each field-caught species were screened for underlying *Orthoflavi*-, *Orthobunya*-, and *Alphavirus* infections using pan-PCRs [19–21] and confirmed negative. All adult mosquitoes were kept at 70% relative humidity, 26°C, and a 12h:12h light:dark photoperiod, including 30 minutes of twilight each. Mosquitoes were fed an 8% fructose solution ad libitum. Two WNV strains were used: WNV strain TOS-09 of L1, isolated in 2008 from a human in Ferrera, Italy (accession no. HM641225, 7^th^ passage, titer: 8.89 × 10^8^ 50% tissue culture infectious dose (TCID_50_) /mL), and WNV strain 1382 of L2, isolated in 2018 from a blackbird in Berlin, Germany (accession no. MH986055, 6^th^ passage, titer: 6.67 × 10^9^ TCID_50_/mL) [22]. Virus propagation and detection of infectious viral particles were performed using Vero cells (*Chlorocebus sabaeus*, ATCC, CCL-81).

### Infection and salivation of mosquitoes

Female mosquitoes (2-15 days old) were deprived of food and water for 24h (*Aedes*) or 48h (*Culex*) prior to an artificial blood meal (50% human blood preservations (type 0), 30% of an 8% fructose solution, 10% fetal bovine serum, 10% virus suspension) for two hours via two 50 µL drops and two saturated cotton swabs per tube. All mosquito species were fed blood meals with 10^5^, 10^6^, and 10^7^ TCID_50_/mL of either WNV lineage, corresponding to viremia levels commonly found in wild birds [15]. *Culex torrentium* was additionally fed with a blood meal containing 10^4^ TCID_50_/mL WNV-L2, as there was still transmission observed at 10^5^ TCID_50_/mL. Fully engorged mosquitoes were incubated at 70% relative humidity, with a 12h:12h light:dark photoperiod, and at 24°C with a ±5°C fluctuation to simulate the natural daily light and temperature change. The temperature extrema were synchronized with the corresponding light and dark phases. An 8% fructose solution was available ad libitum. After 14 days, a salivation assay was performed as previously described [23]. Briefly, wings and legs were removed, and the proboscis was inserted into a capillary tube for 30 min. The saliva solution was placed on Vero cells and incubated (37°C; 5% CO_2_) for seven days. For each experimental condition (viral lineage, mosquito species, blood meal titer), the aim was to analyze 30 mosquito specimens, allowing the identification of transmission efficiencies (TE) of ≥3.33%.

### Downstream processing and detection of WNV

Mosquito bodies, excluding wings and legs, were homogenized individually in 500 µL medium and subjected to RNA extraction using the MagMAX CORE Nucleic Acid Purification Kit (Applied Biosystems; USA). If cells showed cytopathic effect, RNA of the supernatant was extracted using the QIAmp Viral RNA Mini Kit (Qiagen; Germany). All RNA samples were analyzed using the RealStar WNV RT-PCR Kit 2.0 (altona Diagnostics; Germany). RNA of the cell supernatants was used to confirm WNV as the causative agent of the cytopathic effect, whereas RNA from the mosquito bodies was used to quantify the viral RNA per mosquito.

### Data analysis

All calculations, visualizations, and model fitting were performed in R version 4.5.1 [24] using RStudio version 2025.09.1+401 [25] with the packages tidyverse [26] and glmmTMB [27,28]. The infection rate (IR) was calculated as the number of WNV-positive bodies divided by the total number of blood-fed mosquitoes per condition. To determine the TE, the number of mosquitoes with WNV-positive saliva was divided by the total number of blood-fed mosquitoes per condition.

To assess the influence of various predictors on the likelihood of positive infection, and on the body titer (viral RNA copies per mosquito), separate hurdle models were fitted for WNV-L1 and WNV-L2. A hurdle model was chosen due to a large proportion of body titers being zero (i.e. no infection), it overcomes this by separating the analysis into two components: (i) the binary component (zero-inflation) models the probability of detecting viral RNA (i.e. positive infection) using logistic regression with binomial distribution, and (ii) the continuous component analyzes the effects on the body titers, which were log_10_-transformed allowing the use of a linear regression with Gaussian distribution. Mosquito species and blood meal titer (log_10_-transformed) were chosen as predictors for both components of the hurdle models. To convert the outputs of the binary components from likelihood of non-infection to likelihood of infection, the results were inverted. Model diagnostics were performed and indicated a good fit for both models. Pearson residual-based dispersion of the continuous component was 1.16 for the WNV-L1 model and 1.57 for the WNV-L2 model, demonstrating mild to moderate overdispersion. Multicollinearity was assessed, with variance inflation factors of the (i) zero-inflated component being 1.04 for the WNV-L1 model and 1.07 for the WNV-L2 model, and of the (ii) continuous component being 1.08 for the WNV-L1 model and 1.02 for the WNV-L2 model, indicating no relevant collinearity among predictors in either model.

To assess the influence of different factors on the likelihood of detecting positive transmission, a generalized linear model (GLM) with a binomial distribution and logit link function was fitted for WNV-L2. Transmission outcome was taken as the binary response variable, and mosquito species, blood meal titer (log_10_-transformed), and body titer (log_10_-transformed) were used as fixed effects. Model diagnostics were performed and indicated a good fit. Pearson residual-based dispersion was 0.92, demonstrating no overdispersion. Multicollinearity was assessed, with variance inflation factors ranging from 1.02 to 1.43, indicating no relevant collinearity among predictors.

For all models, *Ae. albopictus* was set as reference species. The value 1 was added to all body titers to allow log_10_-transformation of all observations. Model output estimates of the logistic regression models (likelihood of transmission, likelihood of infection) were exponentiated to obtain odds ratios, and 95% confidence intervals were calculated to allow better interpretation of the results.

## Results

### Feeding and survival rates

Feeding rates (fully engorged females per females offered a blood meal) ranged from 22.5% to 47.4% (Supplementary Table 1). Mean survival rates (alive females 14 days post infection per fed females) were generally higher in the two *Culex* species (*Cx. pipiens* biotype *pipiens*: 94.07%, *Cx. torrentium*: 93.30%) compared to *Ae. albopictus* (74.60%) (Supplementary Table 1).

### Infection rates, transmission efficiencies, and minimum infectious doses

All three investigated mosquito species were susceptible to WNV-L1 and L2 infections at all tested blood meal titers (Table 1). For WNV-L1, IRs in *Ae. albopictus* consistently increased with rising blood meal titers, ranging from 46.7% at a blood meal titer of 10^5^ TCID_50_/mL to 60% at 10^7^ TCID_50_/mL, while no virus was detected in the saliva at any blood meal titer. In *Cx. pipiens* biotype *pipiens*, IRs rose from 10.0% to 33.3% at 10^5^ versus 10^7^ TCID_50_/mL, and transmission was only observed at the highest blood meal titer (10^7^ TCID_50_/mL), with a TE of 6.7%. For *Cx. torrentium*, 10^5^ TCID_50_/mL resulted in a slightly higher IR (16.7%) than 10^6^ TCID_50_/mL (13.3%). However, again, the highest blood meal titer of 10^7^ TCID_50_/mL produced the highest IR of 63.3%. Similar to *Cx. pipiens* biotype *pipiens*, WNV was detected in the saliva at the highest blood meal titer only, with an efficiency of 6.7%. Overall, increasing blood meal titers generally resulted in increased IR, and positive saliva was observed only at the highest blood meal titer in *Cx. pipiens* biotype *pipiens* and *Cx. torrentium*, but not in *Ae. albopictus*. For WNV-L2, IRs in *Ae. albopictus* increased from 30.0% at 10^5^ TCID_50_/mL to 86.7% at 10^7^ TCID_50_/mL, with virus detected in the saliva at 10^6^ and 10^7^ TCID_50_/mL (TE = 6.7% and 33.3%). In *Cx. pipiens* biotype *pipiens*, IRs rose from 24.1% to 60.0% with increasing blood meal titers, and transmission was detected at 10^5^ (3.4%) and 10^7^ TCID_50_/mL (10.0%), but not at 10^6^ TCID_50_/mL.

**Table 1:** The number of mosquitoes subjected to salivation assay, the IR (positive bodies/fed mosquitoes), and the TE (positive salivas/fed mosquitoes), stratified by WNV lineage, mosquito species, and blood meal titer. Blood meal titers marked in **bold** indicate the MID necessary for transmission. Mosquitoes were incubated for 14 days at 24±5°C.

| WNV lineage | Mosquito species | Blood meal titer as TCID<sub>50</sub>/mL | No. of tested mosquitoes | Infection rate (no. of mosquitoes with positive body) | Transmission efficiency (no. of mosquitoes with positive saliva) |
| --- | --- | --- | --- | --- | --- |
| Lineage 1 | *Ae. albopictus* | 10<sup>5</sup> | 30 | 46.7% (14) | 0.0% (0) |
|  |  | 10<sup>6</sup> | 30 | 50.0% (15) | 0.0% (0) |
|  |  | 10<sup>7</sup> | 30 | 60.0% (18) | 0.0% (0) |
|  | *Cx. pipiens* biotype *pipiens* | 10<sup>5</sup> | 30 | 10.0% (3) | 0.0% (0) |
|  |  | 10<sup>6</sup> | 30 | 16.7% (5) | 0.0% (0) |
|  |  | **10<sup>7</sup>** | 30 | 33.3% (10) | 6.7% (2) |
|  | *Cx. torrentium* | 10<sup>5</sup> | 30 | 16.7% (5) | 0.0% (0) |
|  |  | 10<sup>6</sup> | 30 | 13.3% (4) | 0.0% (0) |
|  |  | **10<sup>7</sup>** | 30 | 63.3% (19) | 6.7% (2) |
| Lineage 2 | *Ae. albopictus* | 10<sup>5</sup> | 30 | 30.0% (9) | 0.0% (0) |
|  |  | **10<sup>6</sup>** | 30 | 76.7% (23) | 6.7% (2) |
|  |  | 10<sup>7</sup> | 30 | 86.7% (26) | 33.3% (10) |
|  | *Cx. pipiens* biotype *pipiens* | **10<sup>5</sup>** | 29 | 24.1% (7) | 3.4% (1) |
|  |  | 10<sup>6</sup> | 30 | 50.0% (15) | 0.0% (0) |
|  |  | 10<sup>7</sup> | 30 | 60.0% (18) | 10.0% (3) |
|  | *Cx. torrentium* | 10<sup>4</sup> | 30 | 20.0% (6) | 0.0% (0) |
|  |  | **10<sup>5</sup>** | 30 | 33.3% (10) | 3.3% (1) |
|  |  | 10<sup>6</sup> | 33 | 69.7% (23) | 6.1% (2) |
|  |  | 10<sup>7</sup> | 30 | 76.7% (23) | 30.0% (9) |

*Culex torrentium* showed a clear dose-response pattern. For this species, the experiment also included a blood meal titer of 10^4^ TCID_50_/mL for WNV-L2. Testing 10^4^ TCID_50_/mL of L2 was not possible for *Cx. pipiens* biotype *pipiens*, which also still transmitted WNV at the lowest tested blood meal titer, due to the limited number of field-caught specimens. IRs in *Cx. torrentium* ranged from 20% at 10^4^ TCID_50_/mL to 76.7% at 10^7^ TCID_50_/mL. Transmission was detected at 10^5^ TCID_50_/mL with an efficiency of 3.3% and went up to 30.0% at 10^7^ TCID_50_/mL.

The minimum infectious dose (MID, minimum blood meal titer required to cause transmission) was lower for all mosquito species infected with WNV-L2 (Table 1). Both *Culex* species had a MID of 10^7^ TCID_50_/mL for L1 and 10^5^ TCID_50_/mL for L2. *Aedes albopictus*’
s MID was 10^6^ TCID_50_/mL for L2, while not even the highest blood meal titer tested was sufficient to cause transmission for L1.

### Predictors of viral infection, transmission, and body titer

A hurdle model was fitted separately for each viral lineage to assess the factors (mosquito species, blood meal titer) influencing infection. In this model, *Cx. pipiens* biotype *pipiens* had significantly lower odds (odds ratios (ORs): L1 = 0.17, L2 = 0.42) of becoming infected with both lineages compared to *Ae. albopictus*, while *Cx. torrentium* showed the same trend, it was only significant for WNV-L1 (OR = 0.32) (Table 2). In contrast, each log_10_-fold increase in blood meal titer more than doubled the odds of infection for both lineages (ORs: L1 = 2.04, L2 = 2.63). Furthermore, the influence of the different predictors on the body titer was assessed. Once an infection was established, all predictors showed a significant positive effect on body titers for both lineages, with estimates ranging from 0.51 to 1.77 (Table 2). Notably, body titers visually cluster into two groups, a low body titer group and a high body titer group (< and >10^6^ viral RNA copies/body) (Figure 1). Except for six *Ae. albopictus* specimens, all mosquitoes with positive saliva (n = 26) belonged to the high body titer group (Figure 1). These six specimens were all infected with WNV-L2. One of them showed a negative infection result, despite a positive saliva.

**Table 2:** Results of two models assessing the influence of various predictors on (i) the likelihood of positive infection (ii) the body titer (viral RNA/mosquito body) for WNV-L1 and -L2, and (iii) the likelihood of positive transmission for WNV-L2. No transmission model could be fitted for WNV-L1 due to the very limited number of positive saliva samples (n = 4). The infection and body titer model were fitted as a two-step hurdle model, while the transmission model was fitted as a generalized linear model. Effect sizes from the logistic regression components of the transmission and infection models are expressed as odds ratios, whereas parameter estimates are reported for the linear component of the body titer model (denoted by *). The reference category was *Ae. albopictus* for mosquito species. “n.a.” = not available, “n.s.” = non-significant, “CI” = confidence interval.

| Model | Predictor | Odds ratio /<br>*Estimate<br>(95% CI) | p-value | Odds ratio /<br>*Estimate<br>(95% CI) | p-value |
| --- | --- | --- | --- | --- | --- |
|  |  | WNV Lineage 1 |  | WNV Lineage 2 |  |
| Positive infection | *Cx. pipiens* biotype *pipiens* | 0.17 (0.08 – 0.36) | < 0.001 | 0.42 (0.18 – 0.70) | 0.003 |
|  | *Cx. torrentium* | 0.32 (0.16 – 0.64) | 0.001 | 0.76 (0.39 – 1.47) | 0.407 (n.s.) |
|  | Log<sub>10</sub>(Blood meal titer) | 2.04 (1.41 – 2.94) | < 0.001 | 2.63 (1.92 – 3.57) | < 0.001 |
| Body titer | *Cx. pipiens* biotype *pipiens* | *1.38 (0.41 – 2.36) | 0.005 | *1.04 (0.12 – 1.97) | 0.027 |
|  | *Cx. torrentium* | *1.77 (0.92 – 2.62) | < 0.001 | *1.24 (0.41 – 2.07) | 0.003 |
|  | Log<sub>10</sub>(Blood meal titer) | *0.51 (0.05– 0.96) | 0.028 | *1.34 (0.91 – 1.77) | < 0.001 |
| Positive transmission | *Cx. pipiens* biotype *pipiens* | n.a. | n.a. | 0.12 (0.02 – 0.59) | 0.009 |
|  | *Cx. torrentium* | n.a. | n.a. | 0.32 (0.08 – 1.24) | 0.099 (n.s.) |
|  | Log<sub>10</sub>(Blood meal titer) | n.a. | n.a. | 2.24 (0.92 – 5.45) | 0.076 (n.s.) |
|  | Log<sub>10</sub>(Body titer) | n.a. | n.a. | 2.02 (1.55 – 2.65) | < 0.001 |

**Figure 1:**
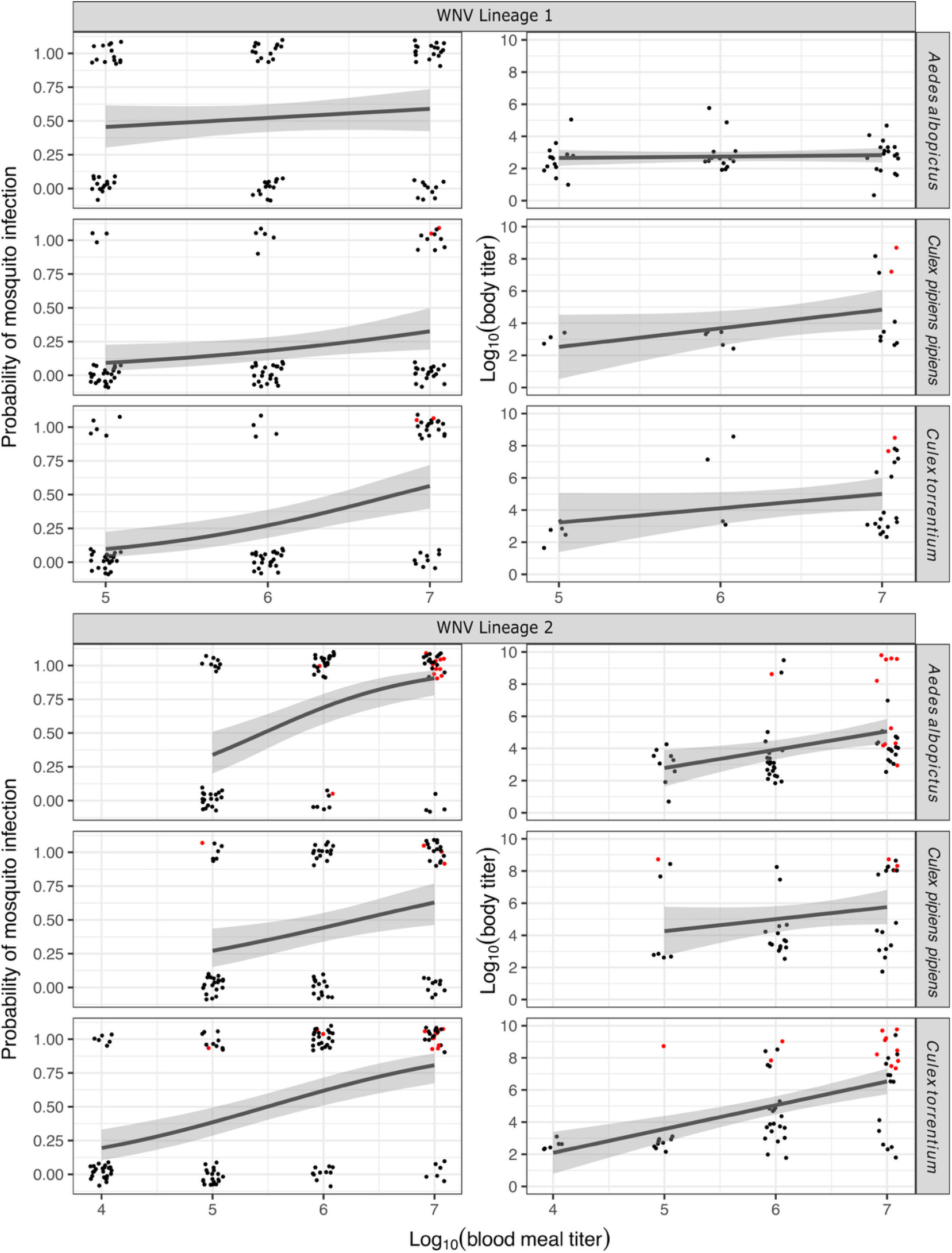
Relationship between the log10-transformed blood meal titer (TCID_50_/mL) and (i) the probability of mosquito infection (left panel, binomial component of the hurdle model) and (ii) the log_10_-transformed body titer (viral RNA/mosquito body) (right panel, linear part of the hurdle model, including only infected mosquitoes with detectable body titers) for *Ae. albopictus, Cx. pipiens* biotype *pipiens*, and *Cx. torrentium* experimentally exposed to WNV-L1 or L2. For (i), logistic regression curves were fitted, whereas for (ii), linear regression curves were fitted. 95% CI are shown in grey. Black dots indicate mosquitoes with a negative saliva, red dots indicate mosquitoes with a positive saliva, jittered for better visibility.

In addition, for WNV-L2, a generalized linear model was fitted to evaluate the predictors of transmission. For WNV-L1, no transmission model could be fitted due to the limited number of positive saliva samples (n = 4), therefore, Figures 2 and 3 only show WNV-L2. In this WNV-L2 model, higher body titers significantly increased the likelihood of detecting positive saliva, with each log_10_-fold increase doubling the odds of transmission (OR = 2.02) (Table 2, Figure 2). Blood meal titer had a similar but non-significant effect (OR = 2.24) (Table 2, Figure 3). *Culex pipiens* biotype *pipiens* significantly reduced the odds of transmission compared to *Ae. albopictus* (OR = 0.12), whereas *Cx. torrentium* also showed reduced odds (OR = 0.32), although this effect was not statistically significant (Table 2). Due to the limited number of positive saliva samples, the same statistical analysis could not be performed for WNV-L1, although it was observed that transmission occurred exclusively at the highest blood meal titer (10^7^ TCID_50_/mL), and only in the two *Culex* species, which showed the same TE of 6.67% (Table 1). However, the proportion of mosquitoes with positive saliva in the higher body titer group (>10^6^ viral RNA copies) is higher in *Cx. pipiens* biotype *pipiens*, compared to *Cx. torrentium* (Figure 1).

**Figure 2:**
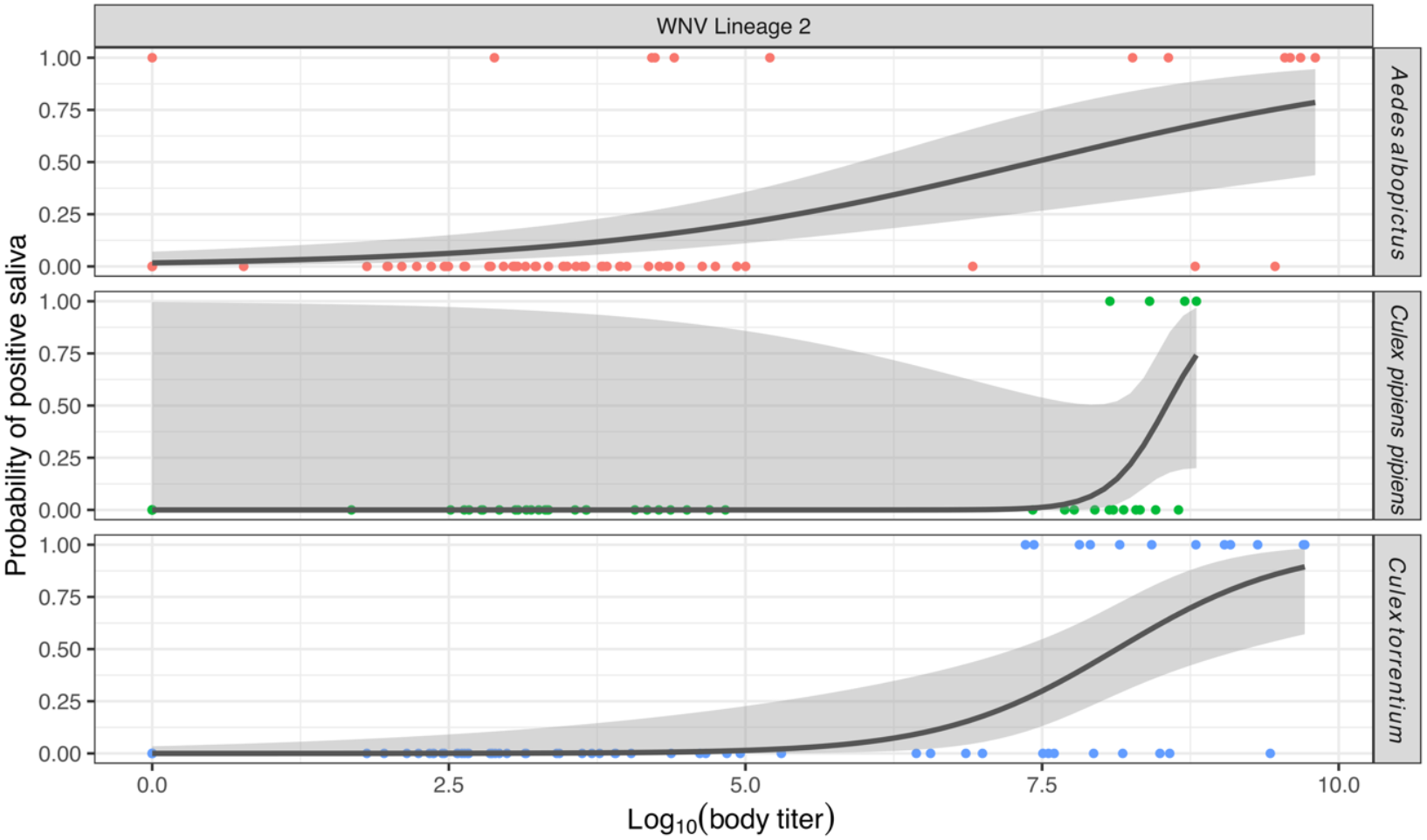
Relationship between the log_10_-transformed body titer (viral RNA/mosquito body) and the probability of positive saliva for *Ae. albopictus, Cx. pipiens* biotype *pipiens*, and *Cx. torrentium* experimentally exposed to WNV-L2. Data points represent observed transmission outcomes (positive = 1; negative = 0), jittered for better visibility. Logistic regression curves with 95% CI in grey were fitted using a binomial generalized linear model.

**Figure 3:**
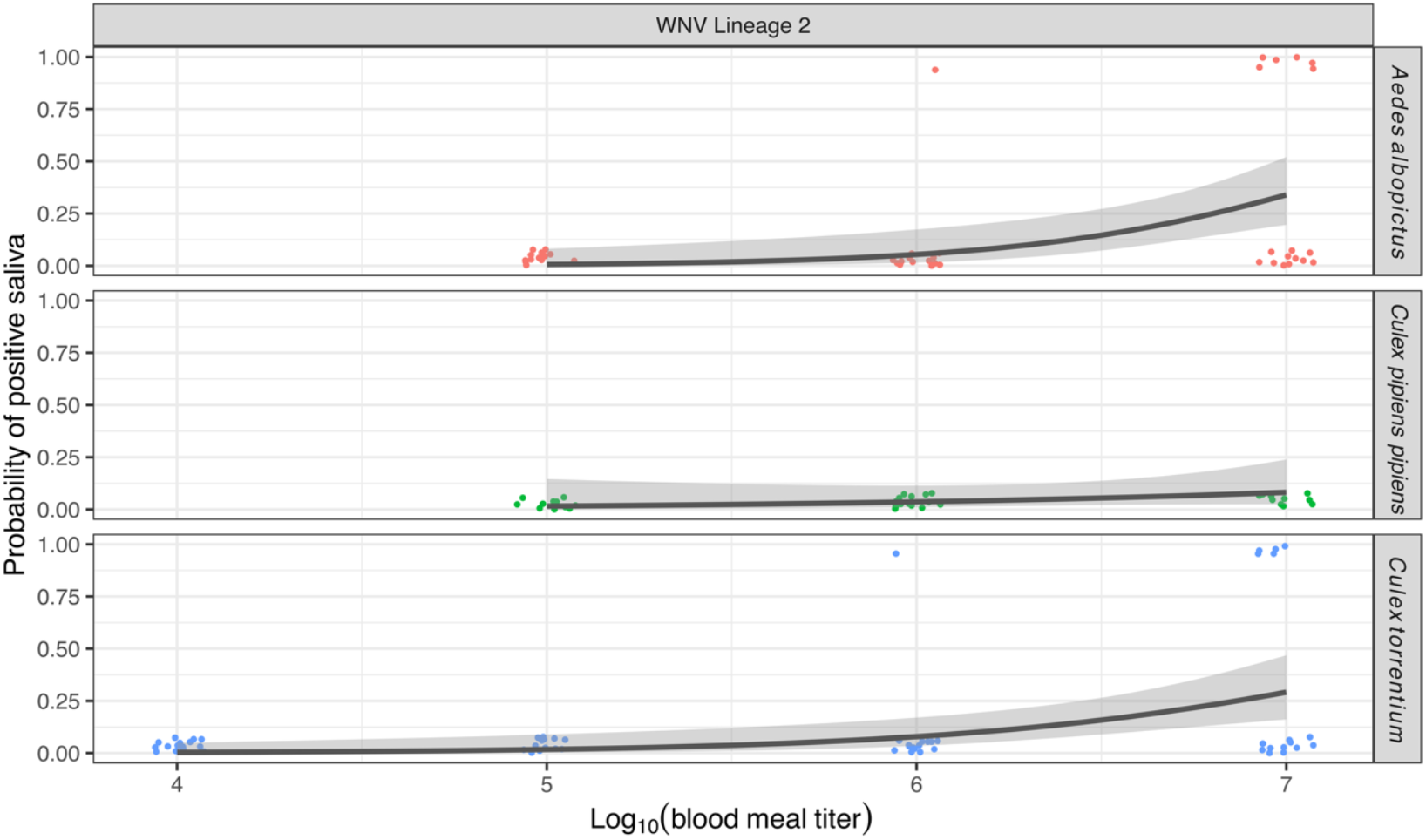
Relationship between the log_10_-transformed blood meal titer (TCID_50_/mL) and the probability of positive saliva for *Ae. albopictus, Cx. pipiens* biotype *pipiens*, and *Cx. torrentium* experimentally exposed to WNV-L2. Data points represent observed transmission outcomes (positive = 1; negative = 0), jittered for better visibility. Logistic regression curves with 95% CI in grey were fitted using a binomial generalized linear model.

## Discussion

Two native European mosquito species, *Cx. pipiens* biotype *pipiens* and *Cx. torrentium*, and the invasive species *Ae. albopictus* were separately infected with WNV-L1 and L2 with blood meal titers ranging from 10^4^ to 10^7^ TCID_50_/mL. All species were susceptible to infection by both lineages, and higher blood meal titers were overall significantly associated with increased IRs and higher body titers. The TEs showed a similar trend, however, differences between blood meal titers were not statistically significant, likely due to the small number of positive saliva samples. This dose-response relationship aligns with previous findings for various mosquito species infected with WNV-L1 [29–31], and has been documented for other arboviruses [32,33].

Beyond the general dose-response effect, there were strong differences both between the mosquito species and the viral lineages. Overall, WNV-L2 resulted in higher IRs, higher TEs, and had lower MIDs across all tested mosquito species compared to L1. Particularly notable was the difference between the two *Culex* species, while both showed the same TEs when infected with WNV-L1, *Cx. torrentium* exhibited a three-times higher TE when infected with WNV-L2 compared to *Cx. pipiens* biotype *pipiens* (30% vs. 10%). A previous study on WNV-L2 in *Cx. pipiens* biotype *pipiens* showed higher TEs of up to 53%, however, a blood meal titer of 10^8^ TCID_50_/mL was used, which was ten-fold higher than the highest one used in our study [34]. Earlier work on WNV-L1 has also identified *Cx. torrentium* as a more competent vector than *Cx. pipiens* biotype *pipiens* with comparable TEs observed in our study [35].

To our knowledge, this study provides the first vector competence assessment of *Cx. torrentium* for WNV-L2. Due to the differences in vector competence between the two tested *Culex* species, it is important to distinguish these morphologically similar species in studies, as it otherwise may obscure key patterns. This is especially important as *Cx. torrentium* is the dominant *Culex* species in Northern Europe, and WNV is predicted to continue its geographic spread towards this area, driven by climate change [2,11].

Although *Ae. albopictus* is not a main WNV vector, it is vector competent for the virus, and WNV has been isolated from field-caught specimens on multiple occasions [12]. Given its rapid expansion across Europe, understanding its potential role as a bridge vector is important [12]. In this study, *Ae. albopictus* was highly susceptible to infection with both WNV lineages. However, *Ae. albopictus* was not competent for WNV-L1, whereas it transmitted WNV-L2 with the highest TEs (max. 33.3%) and at significantly higher odds than *Cx. pipiens* biotype *pipiens*. This contrasts with other European studies that reported transmission of WNV-L1 by *Ae. albopictus* [36]. However, Martinet et al. [36] used a higher mean temperature (28°C) than in this study (24°C), limiting the comparability of results.

Independent of the tested species, higher body titers significantly increased the odds of transmission for WNV-L2, and all mosquitoes that transmitted WNV-L1 exhibited a high body titer. In addition, we also observed a clustering of mosquitoes into high and low body titer groups. Large differences in mosquito body titers have also been observed for other arboviruses, such as Sindbis virus, and have potentially been linked to differential gene regulation and variation in viral population structures [37,38]. However, the underlying causes remain incompletely understood and require detailed investigation. In this study, we also determined the MID for each lineage-mosquito-combination. A low MID can not only expand the range of avian hosts [15] but may also broaden the transmission window, allowing mosquitoes to become infected not only at peak viremia, but also during its rise and decline in amplifying hosts. This extended window may enhance WNV transmission and thereby increase the risk of spillover to humans. Remarkably, the MIDs showed a clear lineage-specific difference. The MID was considerably lower for WNV-L2, with both *Culex* species requiring a 100-fold lower dose to transmit L2 (10^5^ TCID_50_/mL) compared to L1 (10^7^ TCID_50_/mL). For *Ae. albopictus*, the gap was even clearer, as transmission was only observed with WNV-L2 (MID = 10^6^ TCID_50_/mL).

Previous findings on the MID of WNV-L2 in *Ae. albopictus* are comparable to ours. Only 10^7^ and 10^5^ TCID_50_/mL as blood meal titers were tested, and the former was reported as MID [39]. More studies are available for WNV-L1, with the lowest observed MID of 10_^6.7^_ PFU/mL in *Ae. albopictus* from France [36]. This is a stark difference from our findings, where a similar infectious dose did not result in transmission. But again, Martinet et al. [36] used a higher incubation temperature than the present study. Studies with *Cx. pipiens* biotype *pipiens* and *Cx. torrentium* use a high blood meal titer (often 10^7^ viral particles/mL) rather than a range of titers, therefore, comparisons are not feasible [35,40,41]. Our data suggest that the combination of WNV-L2 and *Cx. torrentium* may be of particular concern, as this species exhibited both the lowest MID and a high TE. Together with increasingly favorable environmental conditions, this could contribute to the spread of WNV-L2 in Europe.

Future work should include multiple viral strains to confirm if differences are truly lineage- or strain-specific, and mosquitoes from different regions to assess the influence of different populations on vector competence and MID. Caution is needed when comparing experimental MIDs to natural viremias, as infection and transmission can differ between live hosts and artificial systems [42]. Additionally, TEs measured via artificial salivation have been shown to potentially underestimate the natural transmission potential [43].

In conclusion, this study provides a comprehensive assessment of the infection dynamics of the tested WNV lineages in native European *Cx. pipiens* biotype *pipiens* and *Cx. torrentium*, as well as the invasive *Ae. albopictus*. Distinct viral lineage- and mosquito species-specific differences were observed. WNV-L2 infection consistently resulted in higher TE and lower MIDs across all mosquito species compared to WNV-L1. *Culex torrentium* stood out as a particularly efficient vector for WNV-L2, highlighting a need for closer monitoring. Overall, this study strengthens our understanding of how viral lineage, mosquito species, and blood meal titer interact.

## Acknowledgements

The authors thank Amrei Mack and Unchana Lange (BNITM) for their excellent technical assistance and help in the arthropod breeding facility. We thank Giada Rossini (University of Bologna, Italy) for providing the WNV TOS-09 isolate.

## Funding

This study is part of the CuliFo3 project (grant number FKZ2819107A22) funded by the German Federal Ministry of Agriculture, Food and Regional Identity through the Federal Office for Agriculture and Food; and the German Federal Ministry of Research, Technology and Space under the project NEED and CIMT (grant number 01Kl2022). FGS received funding of the Klaus Tschira Boost Fund, a joint initiative of GSO – Guidance, Skills & Opportunities e.V. and Klaus Tschira Stiftung.

## Author Contributions

Conceptualization: SJ, AH; Methodology: PH; Formal analysis: PH, FGS; Original draft preparation: PH; Review and editing: all authors; Funding acquisition: JSC, AH, RL; Resources: RL, NB; Supervision: AH, SJ; All authors have read and agreed to the published version of the manuscript.

## Competing Interests

The authors declare no competing interests.

## Data availability

The data supporting the findings of this study are available within the article and its supplementary materials. Supplementary Table 1 shows feeding and survival rates. Supplementary Table 2 shows all individual mosquitoes tested, their infection and transmission statuses, and their body titers.

## Notes

### Competing Interest Statement

The authors have declared no competing interest.

