## Supplementary Table 1 for "Not all West Nile virus lineages behave alike: vector competence and minimum infectious dose differences of lineages 1 and 2"

Supplementary Table 1: The number of female mosquitoes offered a blood meal, feeding rate (fully engorged/total), and survival rate (alive after 14 days post infection/fully engorged), stratified by WNV lineage, mosquito species, and blood meal titer. Experiments were conducted at an incubation temperature of 24±5°C.

| **West Nile virus lineage** | **Mosquito species** | **Blood meal titer as TCID_50_/mL** | **Total no. of mosquitoes offered a blood meal** | **Feeding rate (no. of fed individuals)** | **Survival rate (no. of alive individuals after incubation period)** |
| --- | --- | --- | --- | --- | --- |
| Lineage 1 | *Ae. albopictus* | 10^5^ 10^6^ 10^7^ | 143 182 129 | 39.2% (56) 31.3% (57) 41.9% (54) | 83.9% (47) 64.9% (37) 85.2% (46) |
|  | *Cx. pipiens* biotype *pipiens* | 10^5^ 10^6^ 10^7^ | 197 160 139 | 22.8% (45) 22.5% (36) 30.2% (42) | 95.6% (43) 94.4% (34) 95.2% (40) |
|  | *Cx. torrentium* | 10^5^ 10^6^ 10^7^ | 118 119 114 | 44.1% (52) 37.8% (45) 47.4% (54) | 96.2% (50) 97.8% (44)  92.6% (50) |
| Lineage 2 | *Ae. albopictus* | 10^5^ 10^6^ 10^7^ | 113 134 143 | 45.1% (51) 37.3% (50) 46.2% (66) | 78.4% (40) 70.0% (35) 65.2% (43) |
|  | *Cx. pipiens* biotype *pipiens* | 10^5^ 10^6^ 10^7^ | 133 113 150 | 25.6% (34) 42.5% (48) 34.0% (51) | 85.3% (29) 97.8% (47) 96.1% (49) |
|  | *Cx. torrentium* | 10^4^ 10^5^ 10^6^ 10^7^ | 120 123 137 193 | 45.0% (54) 34.1% (42) 26.3% (36) 25.9 % (50) | 94.4% (51) 85.7% (36) 94.4% (34) 92.0% (46) |
