## Supplementary Table 2 for "Not all West Nile virus lineages behave alike: vector competence and minimum infectious dose differences of lineages 1 and 2"

Supplementary Table 2: All individual tested and salivated mosquitoes with their corresponding infection status, transmission status, and body titer. Experiments were conducted at an incubation temperature of 24±5°C.

| **Mosquito Species** | **Virus** | **Blood meal titer in TCID50/mL** | **Result transmission** | **Result infection** | **Mosquito body Cq value** | **Body titer in viral RNA copies / mosquito body** |
| --- | --- | --- | --- | --- | --- | --- |
| Aedes albopictus | WNV Lineage 2 | 1.00E+07 | Neg | Pos | 29.12 | 1.00E+05 |
| Aedes albopictus | WNV Lineage 2 | 1.00E+07 | Neg | Pos | 35.56 | 1.15E+03 |
| Aedes albopictus | WNV Lineage 2 | 1.00E+07 | Neg | Neg |  | 0.00E+00 |
| Aedes albopictus | WNV Lineage 2 | 1.00E+07 | Pos | Pos | 31.66 | 1.73E+04 |
| Aedes albopictus | WNV Lineage 2 | 1.00E+07 | Pos | Pos | 31.12 | 2.51E+04 |
| Aedes albopictus | WNV Lineage 2 | 1.00E+07 | Pos | Pos | 13.91 | 3.90E+09 |
| Aedes albopictus | WNV Lineage 2 | 1.00E+07 | Neg | Pos | 31.28 | 2.25E+04 |
| Aedes albopictus | WNV Lineage 2 | 1.00E+07 | Neg | Pos | 22.78 | 8.24E+06 |
| Aedes albopictus | WNV Lineage 2 | 1.00E+07 | Neg | Pos | 29.97 | 5.58E+04 |
| Aedes albopictus | WNV Lineage 2 | 1.00E+07 | Neg | Pos | 35.01 | 1.69E+03 |
| Aedes albopictus | WNV Lineage 2 | 1.00E+07 | Neg | Neg |  | 0.00E+00 |
| Aedes albopictus | WNV Lineage 2 | 1.00E+07 | Neg | Pos | 37.49 | 2.99E+02 |
| Aedes albopictus | WNV Lineage 2 | 1.00E+07 | Pos | Pos | 36.14 | 7.65E+02 |
| Aedes albopictus | WNV Lineage 2 | 1.00E+07 | Pos | Pos | 14.06 | 3.51E+09 |
| Aedes albopictus | WNV Lineage 2 | 1.00E+07 | Pos | Pos | 28.45 | 1.61E+05 |
| Aedes albopictus | WNV Lineage 2 | 1.00E+07 | Neg | Pos | 32.64 | 8.73E+03 |
| Aedes albopictus | WNV Lineage 2 | 1.00E+07 | Neg | Pos | 31.32 | 2.18E+04 |
| Aedes albopictus | WNV Lineage 2 | 1.00E+07 | Neg | Pos | 31.84 | 1.53E+04 |
| Aedes albopictus | WNV Lineage 2 | 1.00E+07 | Pos | Pos | 13.21 | 6.35E+09 |
| Aedes albopictus | WNV Lineage 2 | 1.00E+07 | Pos | Pos | 18.32 | 1.83E+08 |
| Aedes albopictus | WNV Lineage 2 | 1.00E+07 | Pos | Pos | 31.75 | 1.62E+04 |
| Aedes albopictus | WNV Lineage 2 | 1.00E+07 | Neg | Pos | 30.34 | 4.32E+04 |
| Aedes albopictus | WNV Lineage 2 | 1.00E+07 | Neg | Pos | 33.6 | 4.48E+03 |
| Aedes albopictus | WNV Lineage 2 | 1.00E+07 | Pos | Pos | 13.62 | 4.77E+09 |
| Aedes albopictus | WNV Lineage 2 | 1.00E+07 | Neg | Pos | 34.97 | 1.72E+03 |
| Aedes albopictus | WNV Lineage 2 | 1.00E+07 | Neg | Neg |  | 0.00E+00 |
| Aedes albopictus | WNV Lineage 2 | 1.00E+07 | Neg | Pos | 32.45 | 9.99E+03 |
| Aedes albopictus | WNV Lineage 2 | 1.00E+07 | Neg | Neg |  | 0.00E+00 |
| Aedes albopictus | WNV Lineage 2 | 1.00E+07 | Neg | Pos | 32.62 | 8.87E+03 |
| Aedes albopictus | WNV Lineage 2 | 1.00E+07 | Neg | Pos | 33.15 | 6.12E+03 |
| Aedes albopictus | WNV Lineage 2 | 1.00E+05 | Neg | Pos | 39.13 | 9.63E+01 |
| Aedes albopictus | WNV Lineage 2 | 1.00E+05 | Neg | Neg |  | 0.00E+00 |
| Aedes albopictus | WNV Lineage 2 | 1.00E+05 | Neg | Neg |  | 0.00E+00 |
| Aedes albopictus | WNV Lineage 2 | 1.00E+05 | Neg | Neg |  | 0.00E+00 |
| Aedes albopictus | WNV Lineage 2 | 1.00E+05 | Neg | Neg |  | 0.00E+00 |
| Aedes albopictus | WNV Lineage 2 | 1.00E+05 | Neg | Neg |  | 0.00E+00 |
| Aedes albopictus | WNV Lineage 2 | 1.00E+05 | Neg | Neg |  | 0.00E+00 |
| Aedes albopictus | WNV Lineage 2 | 1.00E+05 | Neg | Neg |  | 0.00E+00 |
| Aedes albopictus | WNV Lineage 2 | 1.00E+05 | Neg | Neg |  | 0.00E+00 |
| Aedes albopictus | WNV Lineage 2 | 1.00E+05 | Neg | Pos | 31.54 | 1.88E+04 |
| Aedes albopictus | WNV Lineage 2 | 1.00E+05 | Neg | Neg |  | 0.00E+00 |
| Aedes albopictus | WNV Lineage 2 | 1.00E+05 | Neg | Pos | 35.89 | 9.14E+02 |
| Aedes albopictus | WNV Lineage 2 | 1.00E+05 | Neg | Pos | 33 | 6.80E+03 |
| Aedes albopictus | WNV Lineage 2 | 1.00E+05 | Neg | Neg |  | 0.00E+00 |
| Aedes albopictus | WNV Lineage 2 | 1.00E+05 | Neg | Neg |  | 0.00E+00 |
| Aedes albopictus | WNV Lineage 2 | 1.00E+05 | Neg | Neg |  | 0.00E+00 |
| Aedes albopictus | WNV Lineage 2 | 1.00E+05 | Neg | Neg |  | 0.00E+00 |
| Aedes albopictus | WNV Lineage 2 | 1.00E+05 | Neg | Neg |  | 0.00E+00 |
| Aedes albopictus | WNV Lineage 2 | 1.00E+05 | Neg | Pos | 37.43 | 3.13E+02 |
| Aedes albopictus | WNV Lineage 2 | 1.00E+05 | Neg | Pos | 34.63 | 2.18E+03 |
| Aedes albopictus | WNV Lineage 2 | 1.00E+05 | Neg | Pos | 43.41 | 4.91E+00 |
| Aedes albopictus | WNV Lineage 2 | 1.00E+05 | Neg | Pos | 34.11 | 3.14E+03 |
| Aedes albopictus | WNV Lineage 2 | 1.00E+05 | Neg | Neg |  | 0.00E+00 |
| Aedes albopictus | WNV Lineage 2 | 1.00E+05 | Neg | Neg |  | 0.00E+00 |
| Aedes albopictus | WNV Lineage 2 | 1.00E+05 | Neg | Neg |  | 0.00E+00 |
| Aedes albopictus | WNV Lineage 2 | 1.00E+05 | Neg | Pos | 33.86 | 3.73E+03 |
| Aedes albopictus | WNV Lineage 2 | 1.00E+05 | Neg | Neg |  | 0.00E+00 |
| Aedes albopictus | WNV Lineage 2 | 1.00E+05 | Neg | Neg |  | 0.00E+00 |
| Aedes albopictus | WNV Lineage 2 | 1.00E+05 | Neg | Neg |  | 0.00E+00 |
| Aedes albopictus | WNV Lineage 2 | 1.00E+05 | Neg | Neg |  | 0.00E+00 |
| Aedes albopictus | WNV Lineage 1 | 1.00E+05 | Neg | Neg |  | 0.00E+00 |
| Aedes albopictus | WNV Lineage 1 | 1.00E+05 | Neg | Pos | 36.75 | 5.00E+02 |
| Aedes albopictus | WNV Lineage 1 | 1.00E+05 | Neg | Neg |  | 0.00E+00 |
| Aedes albopictus | WNV Lineage 1 | 1.00E+05 | Neg | Neg |  | 0.00E+00 |
| Aedes albopictus | WNV Lineage 1 | 1.00E+05 | Neg | Neg |  | 0.00E+00 |
| Aedes albopictus | WNV Lineage 1 | 1.00E+05 | Neg | Neg |  | 0.00E+00 |
| Aedes albopictus | WNV Lineage 1 | 1.00E+05 | Neg | Neg |  | 0.00E+00 |
| Aedes albopictus | WNV Lineage 1 | 1.00E+05 | Neg | Pos | 42.4 | 9.90E+00 |
| Aedes albopictus | WNV Lineage 1 | 1.00E+05 | Neg | Neg |  | 0.00E+00 |
| Aedes albopictus | WNV Lineage 1 | 1.00E+05 | Neg | Neg |  | 0.00E+00 |
| Aedes albopictus | WNV Lineage 1 | 1.00E+05 | Neg | Pos | 34.06 | 3.24E+03 |
| Aedes albopictus | WNV Lineage 1 | 1.00E+05 | Neg | Neg |  | 0.00E+00 |
| Aedes albopictus | WNV Lineage 1 | 1.00E+05 | Neg | Neg |  | 0.00E+00 |
| Aedes albopictus | WNV Lineage 1 | 1.00E+05 | Neg | Neg |  | 0.00E+00 |
| Aedes albopictus | WNV Lineage 1 | 1.00E+05 | Neg | Pos | 28.68 | 1.37E+05 |
| Aedes albopictus | WNV Lineage 1 | 1.00E+05 | Neg | Neg |  | 0.00E+00 |
| Aedes albopictus | WNV Lineage 1 | 1.00E+05 | Neg | Neg |  | 0.00E+00 |
| Aedes albopictus | WNV Lineage 1 | 1.00E+05 | Neg | Neg |  | 0.00E+00 |
| Aedes albopictus | WNV Lineage 1 | 1.00E+05 | Neg | Pos | 36.41 | 6.35E+02 |
| Aedes albopictus | WNV Lineage 1 | 1.00E+05 | Neg | Pos | 39.71 | 6.44E+01 |
| Aedes albopictus | WNV Lineage 1 | 1.00E+07 | Neg | Neg |  | 0.00E+00 |
| Aedes albopictus | WNV Lineage 1 | 1.00E+07 | Neg | Neg |  | 0.00E+00 |
| Aedes albopictus | WNV Lineage 1 | 1.00E+07 | Neg | Neg |  | 0.00E+00 |
| Aedes albopictus | WNV Lineage 1 | 1.00E+07 | Neg | Neg |  | 0.00E+00 |
| Aedes albopictus | WNV Lineage 1 | 1.00E+07 | Neg | Neg |  | 0.00E+00 |
| Aedes albopictus | WNV Lineage 1 | 1.00E+07 | Neg | Neg |  | 0.00E+00 |
| Aedes albopictus | WNV Lineage 1 | 1.00E+07 | Neg | Neg |  | 0.00E+00 |
| Aedes albopictus | WNV Lineage 1 | 1.00E+07 | Neg | Neg |  | 0.00E+00 |
| Aedes albopictus | WNV Lineage 1 | 1.00E+07 | Neg | Pos | 33.07 | 6.44E+03 |
| Aedes albopictus | WNV Lineage 1 | 1.00E+07 | Neg | Pos | 36.46 | 6.12E+02 |
| Aedes albopictus | WNV Lineage 1 | 1.00E+07 | Neg | Pos | 32.54 | 9.59E+02 |
| Aedes albopictus | WNV Lineage 1 | 1.00E+07 | Neg | Pos | 35.83 | 8.69E+01 |
| Aedes albopictus | WNV Lineage 1 | 1.00E+07 | Neg | Pos | 32.99 | 6.93E+02 |
| Aedes albopictus | WNV Lineage 1 | 1.00E+07 | Neg | Neg |  | 0.00E+00 |
| Aedes albopictus | WNV Lineage 1 | 1.00E+07 | Neg | Neg |  | 0.00E+00 |
| Aedes albopictus | WNV Lineage 1 | 1.00E+07 | Neg | Pos | 31.75 | 1.71E+03 |
| Aedes albopictus | WNV Lineage 1 | 1.00E+07 | Neg | Pos | 31.4 | 2.21E+03 |
| Aedes albopictus | WNV Lineage 1 | 1.00E+07 | Neg | Pos | 32.25 | 1.19E+03 |
| Aedes albopictus | WNV Lineage 1 | 1.00E+07 | Neg | Neg |  | 0.00E+00 |
| Aedes albopictus | WNV Lineage 1 | 1.00E+07 | Neg | Neg |  | 0.00E+00 |
| Aedes albopictus | WNV Lineage 1 | 1.00E+07 | Neg | Pos | 28.85 | 1.42E+04 |
| Aedes albopictus | WNV Lineage 1 | 1.00E+07 | Neg | Pos | 36.63 | 4.82E+01 |
| Aedes albopictus | WNV Lineage 1 | 1.00E+07 | Neg | Pos | 41.46 | 1.41E+00 |
| Aedes albopictus | WNV Lineage 1 | 1.00E+07 | Neg | Pos | 33.78 | 3.88E+02 |
| Aedes albopictus | WNV Lineage 1 | 1.00E+07 | Neg | Pos | 35.72 | 9.41E+01 |
| Aedes albopictus | WNV Lineage 1 | 1.00E+07 | Neg | Pos | 37.17 | 3.27E+01 |
| Aedes albopictus | WNV Lineage 1 | 1.00E+07 | Neg | Pos | 31.62 | 1.88E+03 |
| Aedes albopictus | WNV Lineage 1 | 1.00E+07 | Neg | Pos | 32.48 | 1.00E+03 |
| Aedes albopictus | WNV Lineage 1 | 1.00E+07 | Neg | Pos | 27.18 | 4.82E+04 |
| Aedes albopictus | WNV Lineage 1 | 1.00E+07 | Neg | Pos | 33.82 | 3.76E+02 |
| Aedes albopictus | WNV Lineage 1 | 1.00E+06 | Neg | Pos | 35.89 | 8.33E+01 |
| Aedes albopictus | WNV Lineage 1 | 1.00E+06 | Neg | Pos | 35.47 | 1.13E+02 |
| Aedes albopictus | WNV Lineage 1 | 1.00E+06 | Neg | Pos | 35.61 | 1.02E+02 |
| Aedes albopictus | WNV Lineage 1 | 1.00E+06 | Neg | Pos | 34.59 | 2.15E+02 |
| Aedes albopictus | WNV Lineage 1 | 1.00E+06 | Neg | Pos | 32.19 | 1.24E+03 |
| Aedes albopictus | WNV Lineage 1 | 1.00E+06 | Neg | Pos | 33.4 | 5.13E+02 |
| Aedes albopictus | WNV Lineage 1 | 1.00E+06 | Neg | Pos | 32.32 | 1.13E+03 |
| Aedes albopictus | WNV Lineage 1 | 1.00E+06 | Neg | Neg |  | 0.00E+00 |
| Aedes albopictus | WNV Lineage 1 | 1.00E+06 | Neg | Pos | 34.1 | 3.08E+02 |
| Aedes albopictus | WNV Lineage 1 | 1.00E+06 | Neg | Neg |  | 0.00E+00 |
| Aedes albopictus | WNV Lineage 1 | 1.00E+06 | Neg | Pos | 26.35 | 8.91E+04 |
| Aedes albopictus | WNV Lineage 1 | 1.00E+06 | Neg | Pos | 33.47 | 4.86E+02 |
| Aedes albopictus | WNV Lineage 1 | 1.00E+06 | Neg | Pos | 34.33 | 2.59E+02 |
| Aedes albopictus | WNV Lineage 1 | 1.00E+06 | Neg | Pos | 33.58 | 4.50E+02 |
| Aedes albopictus | WNV Lineage 1 | 1.00E+05 | Neg | Neg |  | 0.00E+00 |
| Aedes albopictus | WNV Lineage 1 | 1.00E+05 | Neg | Neg |  | 0.00E+00 |
| Aedes albopictus | WNV Lineage 1 | 1.00E+05 | Neg | Pos | 33.06 | 6.57E+02 |
| Aedes albopictus | WNV Lineage 1 | 1.00E+05 | Neg | Pos | 31.84 | 1.60E+03 |
| Aedes albopictus | WNV Lineage 1 | 1.00E+05 | Neg | Pos | 33.61 | 4.38E+02 |
| Aedes albopictus | WNV Lineage 1 | 1.00E+05 | Neg | Pos | 33.71 | 4.09E+02 |
| Aedes albopictus | WNV Lineage 1 | 1.00E+05 | Neg | Pos | 35.49 | 1.11E+02 |
| Aedes albopictus | WNV Lineage 1 | 1.00E+05 | Neg | Pos | 37.87 | 1.96E+01 |
| Aedes albopictus | WNV Lineage 1 | 1.00E+05 | Neg | Pos | 35.17 | 1.40E+02 |
| Aedes albopictus | WNV Lineage 1 | 1.00E+05 | Neg | Pos | 34.53 | 2.24E+02 |
| Aedes albopictus | WNV Lineage 2 | 1.00E+06 | Pos | Neg |  | 0.00E+00 |
| Aedes albopictus | WNV Lineage 2 | 1.00E+06 | Neg | Pos | 36.26 | 6.35E+01 |
| Aedes albopictus | WNV Lineage 2 | 1.00E+06 | Neg | Pos | 35.71 | 9.45E+01 |
| Aedes albopictus | WNV Lineage 2 | 1.00E+06 | Neg | Pos | 34.18 | 2.89E+02 |
| Aedes albopictus | WNV Lineage 2 | 1.00E+06 | Pos | Pos | 14.96 | 3.67E+08 |
| Aedes albopictus | WNV Lineage 2 | 1.00E+06 | Neg | Pos | 31.01 | 2.95E+03 |
| Aedes albopictus | WNV Lineage 2 | 1.00E+06 | Neg | Pos | 32.03 | 1.40E+03 |
| Aedes albopictus | WNV Lineage 2 | 1.00E+06 | Neg | Neg |  | 0.00E+00 |
| Aedes albopictus | WNV Lineage 2 | 1.00E+06 | Neg | Pos | 32.99 | 6.89E+02 |
| Aedes albopictus | WNV Lineage 2 | 1.00E+06 | Neg | Pos | 32.24 | 1.20E+03 |
| Aedes albopictus | WNV Lineage 2 | 1.00E+06 | Neg | Neg |  | 0.00E+00 |
| Aedes albopictus | WNV Lineage 2 | 1.00E+06 | Neg | Neg |  | 0.00E+00 |
| Aedes albopictus | WNV Lineage 2 | 1.00E+06 | Neg | Pos | 33.62 | 4.36E+02 |
| Aedes albopictus | WNV Lineage 2 | 1.00E+06 | Neg | Pos | 35.32 | 1.26E+02 |
| Aedes albopictus | WNV Lineage 2 | 1.00E+06 | Neg | Pos | 26.41 | 8.46E+04 |
| Aedes albopictus | WNV Lineage 2 | 1.00E+06 | Neg | Pos | 35.35 | 1.10E+03 |
| Aedes albopictus | WNV Lineage 2 | 1.00E+06 | Neg | Pos | 37.93 | 1.69E+02 |
| Aedes albopictus | WNV Lineage 2 | 1.00E+06 | Neg | Pos | 35.32 | 1.12E+03 |
| Aedes albopictus | WNV Lineage 2 | 1.00E+06 | Neg | Neg |  | 0.00E+00 |
| Aedes albopictus | WNV Lineage 2 | 1.00E+06 | Neg | Pos | 14.97 | 2.91E+09 |
| Aedes albopictus | WNV Lineage 2 | 1.00E+06 | Neg | Pos | 30.88 | 2.81E+04 |
| Aedes albopictus | WNV Lineage 2 | 1.00E+06 | Neg | Pos | 35.93 | 7.16E+02 |
| Aedes albopictus | WNV Lineage 2 | 1.00E+06 | Neg | Pos | 32.94 | 6.30E+03 |
| Aedes albopictus | WNV Lineage 2 | 1.00E+06 | Neg | Pos | 34.01 | 2.90E+03 |
| Aedes albopictus | WNV Lineage 2 | 1.00E+06 | Neg | Neg |  | 0.00E+00 |
| Aedes albopictus | WNV Lineage 2 | 1.00E+06 | Neg | Neg |  | 0.00E+00 |
| Aedes albopictus | WNV Lineage 2 | 1.00E+06 | Neg | Pos | 37.54 | 2.24E+02 |
| Aedes albopictus | WNV Lineage 2 | 1.00E+06 | Neg | Pos | 33.48 | 4.24E+03 |
| Aedes albopictus | WNV Lineage 2 | 1.00E+06 | Neg | Pos | 36.65 | 4.24E+02 |
| Aedes albopictus | WNV Lineage 2 | 1.00E+06 | Neg | Pos | 17.11 | 6.17E+08 |
| Culex torrentium | WNV Lineage 2 | 1.00E+07 | Pos | Pos | 14.18 | 5.18E+09 |
| Culex torrentium | WNV Lineage 2 | 1.00E+07 | Neg | Pos | 24.18 | 3.63E+06 |
| Culex torrentium | WNV Lineage 2 | 1.00E+07 | Pos | Pos | 16.32 | 1.09E+09 |
| Culex torrentium | WNV Lineage 2 | 1.00E+07 | Neg | Neg |  | 0.00E+00 |
| Culex torrentium | WNV Lineage 2 | 1.00E+07 | Neg | Neg |  | 0.00E+00 |
| Culex torrentium | WNV Lineage 2 | 1.00E+07 | Neg | Pos | 15.1 | 2.65E+09 |
| Culex torrentium | WNV Lineage 2 | 1.00E+07 | Pos | Pos | 14.22 | 5.04E+09 |
| Culex torrentium | WNV Lineage 2 | 1.00E+07 | Pos | Pos | 19.13 | 1.43E+08 |
| Culex torrentium | WNV Lineage 2 | 1.00E+07 | Neg | Pos | 37.88 | 1.75E+02 |
| Culex torrentium | WNV Lineage 2 | 1.00E+07 | Neg | Pos | 39.28 | 6.35E+01 |
| Culex torrentium | WNV Lineage 2 | 1.00E+07 | Neg | Neg |  | 0.00E+00 |
| Culex torrentium | WNV Lineage 2 | 1.00E+07 | Neg | Pos | 19.82 | 8.60E+07 |
| Culex torrentium | WNV Lineage 2 | 1.00E+07 | Neg | Neg |  | 0.00E+00 |
| Culex torrentium | WNV Lineage 2 | 1.00E+07 | Neg | Neg |  | 0.00E+00 |
| Culex torrentium | WNV Lineage 2 | 1.00E+07 | Neg | Pos | 20.88 | 4.00E+07 |
| Culex torrentium | WNV Lineage 2 | 1.00E+07 | Neg | Pos | 19.04 | 1.51E+08 |
| Culex torrentium | WNV Lineage 2 | 1.00E+07 | Neg | Neg |  | 0.00E+00 |
| Culex torrentium | WNV Lineage 2 | 1.00E+07 | Pos | Pos | 15.44 | 2.07E+09 |
| Culex pipiens biotype pipiens | WNV Lineage 2 | 1.00E+07 | Neg | Neg |  | 0.00E+00 |
| Culex pipiens biotype pipiens | WNV Lineage 2 | 1.00E+07 | Neg | Neg |  | 0.00E+00 |
| Culex pipiens biotype pipiens | WNV Lineage 2 | 1.00E+07 | Neg | Neg |  | 0.00E+00 |
| Culex pipiens biotype pipiens | WNV Lineage 2 | 1.00E+07 | Neg | Neg |  | 0.00E+00 |
| Culex pipiens biotype pipiens | WNV Lineage 2 | 1.00E+07 | Neg | Neg |  | 0.00E+00 |
| Culex pipiens biotype pipiens | WNV Lineage 2 | 1.00E+07 | Neg | Neg |  | 0.00E+00 |
| Culex pipiens biotype pipiens | WNV Lineage 2 | 1.00E+07 | Neg | Pos | 31.45 | 1.86E+04 |
| Culex pipiens biotype pipiens | WNV Lineage 2 | 1.00E+07 | Neg | Pos | 19.41 | 1.17E+08 |
| Culex pipiens biotype pipiens | WNV Lineage 2 | 1.00E+07 | Neg | Neg |  | 0.00E+00 |
| Culex pipiens biotype pipiens | WNV Lineage 2 | 1.00E+07 | Neg | Pos | 36.65 | 4.26E+02 |
| Aedes albopictus | WNV Lineage 1 | 1.00E+06 | Neg | Neg |  | 0.00E+00 |
| Aedes albopictus | WNV Lineage 1 | 1.00E+06 | Neg | Neg |  | 0.00E+00 |
| Aedes albopictus | WNV Lineage 1 | 1.00E+06 | Neg | Neg |  | 0.00E+00 |
| Aedes albopictus | WNV Lineage 1 | 1.00E+06 | Neg | Neg |  | 0.00E+00 |
| Aedes albopictus | WNV Lineage 1 | 1.00E+06 | Neg | Pos | 27.73 | 5.33E+05 |
| Aedes albopictus | WNV Lineage 1 | 1.00E+06 | Neg | Neg |  | 0.00E+00 |
| Aedes albopictus | WNV Lineage 1 | 1.00E+06 | Neg | Neg |  | 0.00E+00 |
| Culex pipiens biotype pipiens | WNV Lineage 2 | 1.00E+07 | Pos | Pos | 19.86 | 1.18E+08 |
| Culex pipiens biotype pipiens | WNV Lineage 2 | 1.00E+07 | Neg | Pos | 19.76 | 1.26E+08 |
| Culex pipiens biotype pipiens | WNV Lineage 2 | 1.00E+07 | Neg | Pos | 32.28 | 2.34E+04 |
| Culex pipiens biotype pipiens | WNV Lineage 2 | 1.00E+07 | Neg | Neg |  | 0.00E+00 |
| Culex pipiens biotype pipiens | WNV Lineage 2 | 1.00E+07 | Neg | Pos | 19.13 | 1.95E+08 |
| Culex pipiens biotype pipiens | WNV Lineage 2 | 1.00E+07 | Neg | Pos | 17.92 | 4.46E+08 |
| Culex pipiens biotype pipiens | WNV Lineage 2 | 1.00E+07 | Neg | Pos | 19.01 | 2.12E+08 |
| Culex pipiens biotype pipiens | WNV Lineage 2 | 1.00E+07 | Neg | Pos | 20.29 | 8.80E+07 |
| Culex pipiens biotype pipiens | WNV Lineage 2 | 1.00E+07 | Pos | Pos | 18.74 | 2.55E+08 |
| Culex pipiens biotype pipiens | WNV Lineage 2 | 1.00E+07 | Neg | Pos | 36.36 | 1.42E+03 |
| Culex pipiens biotype pipiens | WNV Lineage 2 | 1.00E+07 | Neg | Pos | 35.83 | 2.05E+03 |
| Culex pipiens biotype pipiens | WNV Lineage 2 | 1.00E+07 | Pos | Pos | 17.42 | 6.32E+08 |
| Culex pipiens biotype pipiens | WNV Lineage 2 | 1.00E+07 | Neg | Pos | 20.87 | 5.90E+07 |
| Culex pipiens biotype pipiens | WNV Lineage 2 | 1.00E+07 | Neg | Pos | 30.73 | 6.78E+04 |
| Culex torrentium | WNV Lineage 2 | 1.00E+06 | Neg | Neg |  | 0.00E+00 |
| Culex torrentium | WNV Lineage 2 | 1.00E+06 | Neg | Neg |  | 0.00E+00 |
| Culex torrentium | WNV Lineage 2 | 1.00E+06 | Neg | Neg |  | 0.00E+00 |
| Culex torrentium | WNV Lineage 2 | 1.00E+06 | Neg | Neg |  | 0.00E+00 |
| Culex torrentium | WNV Lineage 2 | 1.00E+06 | Neg | Neg |  | 0.00E+00 |
| Culex torrentium | WNV Lineage 2 | 1.00E+06 | Neg | Pos | 29.15 | 2.01E+05 |
| Culex torrentium | WNV Lineage 2 | 1.00E+06 | Neg | Neg |  | 0.00E+00 |
| Culex pipiens biotype pipiens | WNV Lineage 2 | 1.00E+06 | Neg | Pos | 34.98 | 3.66E+03 |
| Culex pipiens biotype pipiens | WNV Lineage 2 | 1.00E+06 | Neg | Neg |  | 0.00E+00 |
| Culex pipiens biotype pipiens | WNV Lineage 2 | 1.00E+06 | Neg | Neg |  | 0.00E+00 |
| Culex torrentium | WNV Lineage 2 | 1.00E+06 | Neg | Pos | 18.17 | 3.76E+08 |
| Culex torrentium | WNV Lineage 2 | 1.00E+06 | Neg | Pos | 34.51 | 5.08E+03 |
| Culex torrentium | WNV Lineage 2 | 1.00E+06 | Neg | Neg |  | 0.00E+00 |
| Culex torrentium | WNV Lineage 2 | 1.00E+06 | Neg | Pos | 34.78 | 4.20E+03 |
| Culex torrentium | WNV Lineage 2 | 1.00E+06 | Neg | Neg |  | 0.00E+00 |
| Culex torrentium | WNV Lineage 2 | 1.00E+06 | Neg | Pos | 33.85 | 7.96E+03 |
| Culex torrentium | WNV Lineage 2 | 1.00E+06 | Neg | Pos | 18.44 | 3.11E+08 |
| Culex pipiens biotype pipiens | WNV Lineage 2 | 1.00E+06 | Neg | Neg |  | 0.00E+00 |
| Culex pipiens biotype pipiens | WNV Lineage 2 | 1.00E+06 | Neg | Neg |  | 0.00E+00 |
| Culex pipiens biotype pipiens | WNV Lineage 2 | 1.00E+06 | Neg | Neg |  | 0.00E+00 |
| Culex pipiens biotype pipiens | WNV Lineage 2 | 1.00E+06 | Neg | Pos | 32.94 | 1.49E+04 |
| Culex pipiens biotype pipiens | WNV Lineage 2 | 1.00E+06 | Neg | Neg |  | 0.00E+00 |
| Culex pipiens biotype pipiens | WNV Lineage 2 | 1.00E+06 | Neg | Pos | 35.73 | 2.19E+03 |
| Culex pipiens biotype pipiens | WNV Lineage 2 | 1.00E+06 | Neg | Pos | 34.67 | 4.53E+03 |
| Culex pipiens biotype pipiens | WNV Lineage 2 | 1.00E+06 | Neg | Neg |  | 0.00E+00 |
| Culex pipiens biotype pipiens | WNV Lineage 2 | 1.00E+06 | Neg | Pos | 34.66 | 4.58E+03 |
| Culex pipiens biotype pipiens | WNV Lineage 2 | 1.00E+06 | Neg | Pos | 36.01 | 1.81E+03 |
| Culex pipiens biotype pipiens | WNV Lineage 2 | 1.00E+06 | Neg | Pos | 36.67 | 1.15E+03 |
| Culex pipiens biotype pipiens | WNV Lineage 2 | 1.00E+06 | Neg | Pos | 31.18 | 4.98E+04 |
| Culex pipiens biotype pipiens | WNV Lineage 2 | 1.00E+06 | Neg | Pos | 31.82 | 3.21E+04 |
| Culex pipiens biotype pipiens | WNV Lineage 2 | 1.00E+06 | Neg | Neg |  | 0.00E+00 |
| Culex pipiens biotype pipiens | WNV Lineage 2 | 1.00E+06 | Neg | Neg |  | 0.00E+00 |
| Culex pipiens biotype pipiens | WNV Lineage 2 | 1.00E+06 | Neg | Pos | 35.78 | 2.12E+03 |
| Culex pipiens biotype pipiens | WNV Lineage 2 | 1.00E+06 | Neg | Pos | 19.47 | 1.54E+08 |
| Culex pipiens biotype pipiens | WNV Lineage 2 | 1.00E+06 | Neg | Neg |  | 0.00E+00 |
| Culex pipiens biotype pipiens | WNV Lineage 2 | 1.00E+06 | Neg | Neg |  | 0.00E+00 |
| Culex pipiens biotype pipiens | WNV Lineage 2 | 1.00E+06 | Neg | Pos | 33.29 | 1.17E+04 |
| Culex torrentium | WNV Lineage 2 | 1.00E+07 | Neg | Pos | 34.37 | 4.55E+02 |
| Culex torrentium | WNV Lineage 2 | 1.00E+07 | Pos | Pos | 16.58 | 2.66E+08 |
| Culex torrentium | WNV Lineage 2 | 1.00E+07 | Pos | Pos | 19.86 | 2.29E+07 |
| Culex torrentium | WNV Lineage 2 | 1.00E+07 | Pos | Pos | 19.64 | 2.70E+07 |
| Culex torrentium | WNV Lineage 2 | 1.00E+07 | Neg | Pos | 20.98 | 9.95E+06 |
| Culex torrentium | WNV Lineage 2 | 1.00E+07 | Neg | Neg |  | 0.00E+00 |
| Culex torrentium | WNV Lineage 2 | 1.00E+07 | Pos | Pos | 18.46 | 6.53E+07 |
| Culex torrentium | WNV Lineage 2 | 1.00E+07 | Neg | Pos | 22.7 | 2.75E+06 |
| Culex torrentium | WNV Lineage 2 | 1.00E+07 | Neg | Pos | 35.27 | 2.33E+02 |
| Culex torrentium | WNV Lineage 2 | 1.00E+07 | Neg | Pos | 21.41 | 7.20E+06 |
| Culex pipiens biotype pipiens | WNV Lineage 2 | 1.00E+07 | Neg | Neg |  | 0.00E+00 |
| Culex pipiens biotype pipiens | WNV Lineage 2 | 1.00E+07 | Neg | Neg |  | 0.00E+00 |
| Culex pipiens biotype pipiens | WNV Lineage 2 | 1.00E+07 | Neg | Pos | 37.41 | 4.68E+01 |
| Culex pipiens biotype pipiens | WNV Lineage 2 | 1.00E+07 | Neg | Neg |  | 0.00E+00 |
| Culex pipiens biotype pipiens | WNV Lineage 2 | 1.00E+07 | Neg | Neg |  | 0.00E+00 |
| Culex pipiens biotype pipiens | WNV Lineage 2 | 1.00E+07 | Neg | Pos | 33.06 | 1.21E+03 |
| Culex pipiens biotype pipiens | WNV Lineage 2 | 1.00E+06 | Neg | Neg |  | 0.00E+00 |
| Culex pipiens biotype pipiens | WNV Lineage 2 | 1.00E+06 | Neg | Neg |  | 0.00E+00 |
| Culex pipiens biotype pipiens | WNV Lineage 2 | 1.00E+06 | Neg | Pos | 32.72 | 1.56E+03 |
| Culex pipiens biotype pipiens | WNV Lineage 2 | 1.00E+06 | Neg | Pos | 34.82 | 3.26E+02 |
| Culex pipiens biotype pipiens | WNV Lineage 2 | 1.00E+06 | Neg | Neg |  | 0.00E+00 |
| Culex pipiens biotype pipiens | WNV Lineage 2 | 1.00E+06 | Neg | Pos | 19.68 | 2.64E+07 |
| Culex pipiens biotype pipiens | WNV Lineage 2 | 1.00E+06 | Neg | Neg |  | 0.00E+00 |
| Culex torrentium | WNV Lineage 2 | 1.00E+06 | Neg | Neg |  | 0.00E+00 |
| Culex torrentium | WNV Lineage 2 | 1.00E+06 | Neg | Pos | 33.73 | 7.34E+02 |
| Culex torrentium | WNV Lineage 2 | 1.00E+06 | Pos | Pos | 18.18 | 8.06E+07 |
| Culex torrentium | WNV Lineage 2 | 1.00E+06 | Neg | Pos | 36.55 | 8.91E+01 |
| Culex torrentium | WNV Lineage 2 | 1.00E+06 | Neg | Neg |  | 0.00E+00 |
| Culex torrentium | WNV Lineage 2 | 1.00E+06 | Neg | Pos | 37.01 | 6.35E+01 |
| Culex torrentium | WNV Lineage 2 | 1.00E+06 | Neg | Pos | 35.92 | 9.77E+02 |
| Culex torrentium | WNV Lineage 2 | 1.00E+06 | Neg | Pos | 33.41 | 5.90E+03 |
| Culex torrentium | WNV Lineage 2 | 1.00E+06 | Pos | Pos | 16.25 | 1.22E+09 |
| Culex torrentium | WNV Lineage 2 | 1.00E+06 | Neg | Pos | 36.13 | 8.42E+02 |
| Culex torrentium | WNV Lineage 2 | 1.00E+06 | Neg | Pos | 29.58 | 9.05E+04 |
| Culex torrentium | WNV Lineage 2 | 1.00E+06 | Neg | Pos | 31.45 | 2.37E+04 |
| Culex torrentium | WNV Lineage 2 | 1.00E+06 | Neg | Pos | 29.94 | 6.98E+04 |
| Culex torrentium | WNV Lineage 2 | 1.00E+06 | Neg | Pos | 34.57 | 2.57E+03 |
| Culex torrentium | WNV Lineage 2 | 1.00E+06 | Neg | Pos | 21.35 | 3.21E+07 |
| Culex torrentium | WNV Lineage 2 | 1.00E+06 | Neg | Pos | 21.2 | 3.58E+07 |
| Culex torrentium | WNV Lineage 2 | 1.00E+06 | Neg | Pos | 30.51 | 4.64E+04 |
| Culex torrentium | WNV Lineage 2 | 1.00E+06 | Neg | Pos | 32.97 | 8.06E+03 |
| Culex torrentium | WNV Lineage 2 | 1.00E+06 | Neg | Pos | 30.68 | 4.13E+04 |
| Culex torrentium | WNV Lineage 2 | 1.00E+07 | Neg | Pos | 32.54 | 1.09E+04 |
| Culex torrentium | WNV Lineage 2 | 1.00E+07 | Neg | Pos | 34.5 | 2.70E+03 |
| Culex torrentium | WNV Lineage 2 | 1.00E+05 | Neg | Pos | 36.39 | 7.02E+02 |
| Culex torrentium | WNV Lineage 2 | 1.00E+05 | Pos | Pos | 17.19 | 6.26E+08 |
| Culex torrentium | WNV Lineage 2 | 1.00E+05 | Neg | Neg |  | 0.00E+00 |
| Culex torrentium | WNV Lineage 2 | 1.00E+05 | Neg | Neg |  | 0.00E+00 |
| Culex torrentium | WNV Lineage 2 | 1.00E+05 | Neg | Neg |  | 0.00E+00 |
| Culex torrentium | WNV Lineage 2 | 1.00E+05 | Neg | Neg |  | 0.00E+00 |
| Culex torrentium | WNV Lineage 2 | 1.00E+05 | Neg | Neg |  | 0.00E+00 |
| Culex torrentium | WNV Lineage 2 | 1.00E+05 | Neg | Pos | 36.35 | 7.20E+02 |
| Culex torrentium | WNV Lineage 2 | 1.00E+05 | Neg | Pos | 35.4 | 1.42E+03 |
| Culex torrentium | WNV Lineage 2 | 1.00E+05 | Neg | Pos | 37.26 | 3.76E+02 |
| Culex torrentium | WNV Lineage 2 | 1.00E+05 | Neg | Neg |  | 0.00E+00 |
| Culex torrentium | WNV Lineage 2 | 1.00E+05 | Neg | Neg |  | 0.00E+00 |
| Culex torrentium | WNV Lineage 2 | 1.00E+05 | Neg | Neg |  | 0.00E+00 |
| Culex torrentium | WNV Lineage 2 | 1.00E+05 | Neg | Neg |  | 0.00E+00 |
| Culex torrentium | WNV Lineage 2 | 1.00E+05 | Neg | Neg |  | 0.00E+00 |
| Culex torrentium | WNV Lineage 2 | 1.00E+05 | Neg | Pos | 37.08 | 4.29E+02 |
| Culex torrentium | WNV Lineage 2 | 1.00E+05 | Neg | Neg |  | 0.00E+00 |
| Culex torrentium | WNV Lineage 2 | 1.00E+05 | Neg | Neg |  | 0.00E+00 |
| Culex torrentium | WNV Lineage 2 | 1.00E+05 | Neg | Neg |  | 0.00E+00 |
| Culex torrentium | WNV Lineage 2 | 1.00E+05 | Neg | Neg |  | 0.00E+00 |
| Culex torrentium | WNV Lineage 2 | 1.00E+05 | Neg | Neg |  | 0.00E+00 |
| Culex torrentium | WNV Lineage 2 | 1.00E+05 | Neg | Pos | 36.59 | 7.79E+02 |
| Culex torrentium | WNV Lineage 2 | 1.00E+05 | Neg | Neg |  | 0.00E+00 |
| Culex torrentium | WNV Lineage 2 | 1.00E+05 | Neg | Pos | 37.99 | 2.78E+02 |
| Culex torrentium | WNV Lineage 2 | 1.00E+05 | Neg | Pos | 38.94 | 1.39E+02 |
| Culex torrentium | WNV Lineage 2 | 1.00E+05 | Neg | Neg |  | 0.00E+00 |
| Culex torrentium | WNV Lineage 2 | 1.00E+05 | Neg | Pos | 37.6 | 3.71E+02 |
| Culex torrentium | WNV Lineage 2 | 1.00E+04 | Neg | Neg |  | 0.00E+00 |
| Culex torrentium | WNV Lineage 2 | 1.00E+04 | Neg | Neg |  | 0.00E+00 |
| Culex torrentium | WNV Lineage 2 | 1.00E+04 | Neg | Pos | 37.2 | 2.16E+02 |
| Culex torrentium | WNV Lineage 2 | 1.00E+04 | Neg | Neg |  | 0.00E+00 |
| Culex torrentium | WNV Lineage 2 | 1.00E+04 | Neg | Neg |  | 0.00E+00 |
| Culex torrentium | WNV Lineage 2 | 1.00E+04 | Neg | Neg |  | 0.00E+00 |
| Culex torrentium | WNV Lineage 2 | 1.00E+04 | Neg | Neg |  | 0.00E+00 |
| Culex torrentium | WNV Lineage 2 | 1.00E+04 | Neg | Neg |  | 0.00E+00 |
| Culex torrentium | WNV Lineage 2 | 1.00E+04 | Neg | Neg |  | 0.00E+00 |
| Culex torrentium | WNV Lineage 2 | 1.00E+04 | Neg | Neg |  | 0.00E+00 |
| Culex torrentium | WNV Lineage 2 | 1.00E+04 | Neg | Neg |  | 0.00E+00 |
| Culex torrentium | WNV Lineage 2 | 1.00E+04 | Neg | Neg |  | 0.00E+00 |
| Culex torrentium | WNV Lineage 2 | 1.00E+04 | Neg | Neg |  | 0.00E+00 |
| Culex pipiens biotype pipiens | WNV Lineage 1 | 1.00E+07 | Neg | Neg |  | 0.00E+00 |
| Culex pipiens biotype pipiens | WNV Lineage 1 | 1.00E+07 | Neg | Pos | 19.87 | 1.59E+08 |
| Culex pipiens biotype pipiens | WNV Lineage 1 | 1.00E+07 | Pos | Pos | 23.08 | 1.30E+07 |
| Culex pipiens biotype pipiens | WNV Lineage 1 | 1.00E+07 | Neg | Neg |  | 0.00E+00 |
| Culex pipiens biotype pipiens | WNV Lineage 1 | 1.00E+07 | Neg | Neg |  | 0.00E+00 |
| Culex pipiens biotype pipiens | WNV Lineage 1 | 1.00E+07 | Neg | Pos | 23.25 | 1.14E+07 |
| Culex pipiens biotype pipiens | WNV Lineage 1 | 1.00E+07 | Neg | Neg |  | 0.00E+00 |
| Culex pipiens biotype pipiens | WNV Lineage 1 | 1.00E+07 | Neg | Neg |  | 0.00E+00 |
| Culex torrentium | WNV Lineage 2 | 1.00E+04 | Neg | Pos | 36.23 | 4.64E+02 |
| Culex torrentium | WNV Lineage 2 | 1.00E+04 | Neg | Pos | 36.39 | 4.06E+02 |
| Culex torrentium | WNV Lineage 2 | 1.00E+04 | Neg | Neg |  | 0.00E+00 |
| Culex torrentium | WNV Lineage 2 | 1.00E+04 | Neg | Pos | 36.84 | 2.88E+02 |
| Culex torrentium | WNV Lineage 2 | 1.00E+04 | Neg | Pos | 34.81 | 1.39E+03 |
| Culex torrentium | WNV Lineage 2 | 1.00E+04 | Neg | Neg |  | 0.00E+00 |
| Culex torrentium | WNV Lineage 2 | 1.00E+04 | Neg | Neg |  | 0.00E+00 |
| Culex torrentium | WNV Lineage 2 | 1.00E+04 | Neg | Neg |  | 0.00E+00 |
| Culex torrentium | WNV Lineage 2 | 1.00E+04 | Neg | Neg |  | 0.00E+00 |
| Culex torrentium | WNV Lineage 2 | 1.00E+04 | Neg | Neg |  | 0.00E+00 |
| Culex torrentium | WNV Lineage 2 | 1.00E+04 | Neg | Pos | 37.1 | 2.35E+02 |
| Culex torrentium | WNV Lineage 2 | 1.00E+04 | Neg | Neg |  | 0.00E+00 |
| Culex torrentium | WNV Lineage 2 | 1.00E+04 | Neg | Neg |  | 0.00E+00 |
| Culex torrentium | WNV Lineage 2 | 1.00E+04 | Neg | Neg |  | 0.00E+00 |
| Culex torrentium | WNV Lineage 2 | 1.00E+04 | Neg | Neg |  | 0.00E+00 |
| Culex torrentium | WNV Lineage 2 | 1.00E+04 | Neg | Neg |  | 0.00E+00 |
| Culex torrentium | WNV Lineage 2 | 1.00E+04 | Neg | Neg |  | 0.00E+00 |
| Culex pipiens biotype pipiens | WNV Lineage 1 | 1.00E+07 | Neg | Pos | 36.41 | 4.01E+02 |
| Culex pipiens biotype pipiens | WNV Lineage 1 | 1.00E+07 | Neg | Neg |  | 0.00E+00 |
| Culex pipiens biotype pipiens | WNV Lineage 1 | 1.00E+07 | Neg | Neg |  | 0.00E+00 |
| Culex pipiens biotype pipiens | WNV Lineage 1 | 1.00E+07 | Pos | Pos | 18.49 | 4.64E+08 |
| Culex torrentium | WNV Lineage 1 | 1.00E+07 | Neg | Pos | 34.68 | 1.54E+03 |
| Culex torrentium | WNV Lineage 1 | 1.00E+07 | Neg | Pos | 25.95 | 1.40E+06 |
| Culex torrentium | WNV Lineage 1 | 1.00E+07 | Neg | Pos | 35.7 | 6.98E+02 |
| Culex torrentium | WNV Lineage 1 | 1.00E+07 | Neg | Pos | 23.74 | 7.79E+06 |
| Culex torrentium | WNV Lineage 1 | 1.00E+07 | Neg | Pos | 33.87 | 2.89E+03 |
| Culex torrentium | WNV Lineage 1 | 1.00E+07 | Neg | Pos | 35.17 | 1.05E+03 |
| Culex torrentium | WNV Lineage 1 | 1.00E+07 | Neg | Pos | 36.05 | 5.31E+02 |
| Culex torrentium | WNV Lineage 1 | 1.00E+07 | Pos | Pos | 19 | 3.13E+08 |
| Culex torrentium | WNV Lineage 1 | 1.00E+07 | Neg | Pos | 32.9 | 6.17E+03 |
| Culex torrentium | WNV Lineage 1 | 1.00E+07 | Neg | Pos | 33.76 | 3.16E+03 |
| Culex torrentium | WNV Lineage 1 | 1.00E+07 | Neg | Pos | 34.4 | 1.93E+03 |
| Culex torrentium | WNV Lineage 1 | 1.00E+07 | Neg | Neg |  | 0.00E+00 |
| Culex torrentium | WNV Lineage 1 | 1.00E+07 | Neg | Neg |  | 0.00E+00 |
| Culex torrentium | WNV Lineage 1 | 1.00E+07 | Neg | Pos | 36.55 | 3.58E+02 |
| Culex torrentium | WNV Lineage 1 | 1.00E+07 | Neg | Pos | 35.17 | 1.05E+03 |
| Culex torrentium | WNV Lineage 1 | 1.00E+06 | Neg | Pos | 34.3 | 2.08E+03 |
| Culex torrentium | WNV Lineage 1 | 1.00E+06 | Neg | Neg |  | 0.00E+00 |
| Culex torrentium | WNV Lineage 1 | 1.00E+06 | Neg | Neg |  | 0.00E+00 |
| Culex torrentium | WNV Lineage 1 | 1.00E+06 | Neg | Neg |  | 0.00E+00 |
| Culex torrentium | WNV Lineage 1 | 1.00E+06 | Neg | Neg |  | 0.00E+00 |
| Culex torrentium | WNV Lineage 1 | 1.00E+06 | Neg | Neg |  | 0.00E+00 |
| Culex torrentium | WNV Lineage 1 | 1.00E+06 | Neg | Neg |  | 0.00E+00 |
| Culex torrentium | WNV Lineage 1 | 1.00E+06 | Neg | Neg |  | 0.00E+00 |
| Culex torrentium | WNV Lineage 1 | 1.00E+06 | Neg | Neg |  | 0.00E+00 |
| Culex torrentium | WNV Lineage 1 | 1.00E+06 | Neg | Neg |  | 0.00E+00 |
| Culex torrentium | WNV Lineage 1 | 1.00E+06 | Neg | Neg |  | 0.00E+00 |
| Culex torrentium | WNV Lineage 1 | 1.00E+06 | Neg | Neg |  | 0.00E+00 |
| Culex torrentium | WNV Lineage 1 | 1.00E+06 | Neg | Pos | 22.95 | 1.44E+07 |
| Culex torrentium | WNV Lineage 1 | 1.00E+06 | Neg | Neg |  | 0.00E+00 |
| Culex torrentium | WNV Lineage 1 | 1.00E+06 | Neg | Pos | 18.71 | 3.94E+08 |
| Culex torrentium | WNV Lineage 1 | 1.00E+05 | Neg | Neg |  | 0.00E+00 |
| Culex torrentium | WNV Lineage 1 | 1.00E+05 | Neg | Neg |  | 0.00E+00 |
| Culex torrentium | WNV Lineage 1 | 1.00E+05 | Neg | Neg |  | 0.00E+00 |
| Culex torrentium | WNV Lineage 1 | 1.00E+05 | Neg | Neg |  | 0.00E+00 |
| Culex torrentium | WNV Lineage 1 | 1.00E+05 | Neg | Neg |  | 0.00E+00 |
| Culex torrentium | WNV Lineage 1 | 1.00E+05 | Neg | Neg |  | 0.00E+00 |
| Culex torrentium | WNV Lineage 1 | 1.00E+05 | Neg | Pos | 35.73 | 6.80E+02 |
| Culex torrentium | WNV Lineage 1 | 1.00E+05 | Neg | Pos | 39.04 | 5.13E+01 |
| Culex torrentium | WNV Lineage 1 | 1.00E+05 | Neg | Neg |  | 0.00E+00 |
| Culex torrentium | WNV Lineage 1 | 1.00E+05 | Neg | Neg |  | 0.00E+00 |
| Culex torrentium | WNV Lineage 1 | 1.00E+05 | Neg | Neg |  | 0.00E+00 |
| Culex torrentium | WNV Lineage 1 | 1.00E+05 | Neg | Neg |  | 0.00E+00 |
| Culex torrentium | WNV Lineage 1 | 1.00E+05 | Neg | Neg |  | 0.00E+00 |
| Culex torrentium | WNV Lineage 1 | 1.00E+05 | Neg | Neg |  | 0.00E+00 |
| Culex torrentium | WNV Lineage 1 | 1.00E+05 | Neg | Neg |  | 0.00E+00 |
| Culex torrentium | WNV Lineage 1 | 1.00E+07 | Neg | Neg |  | 0.00E+00 |
| Culex torrentium | WNV Lineage 1 | 1.00E+07 | Neg | Neg |  | 0.00E+00 |
| Culex torrentium | WNV Lineage 1 | 1.00E+07 | Neg | Pos | 25.17 | 2.55E+06 |
| Culex torrentium | WNV Lineage 1 | 1.00E+07 | Neg | Pos | 21.27 | 5.36E+07 |
| Culex torrentium | WNV Lineage 1 | 1.00E+07 | Neg | Pos | 20.03 | 7.25E+07 |
| Culex torrentium | WNV Lineage 1 | 1.00E+07 | Neg | Neg |  | 0.00E+00 |
| Culex torrentium | WNV Lineage 1 | 1.00E+07 | Neg | Neg |  | 0.00E+00 |
| Culex torrentium | WNV Lineage 1 | 1.00E+07 | Neg | Neg |  | 0.00E+00 |
| Culex torrentium | WNV Lineage 1 | 1.00E+07 | Pos | Pos | 20.94 | 3.92E+07 |
| Culex torrentium | WNV Lineage 1 | 1.00E+07 | Neg | Pos | 38.65 | 2.61E+02 |
| Culex torrentium | WNV Lineage 1 | 1.00E+07 | Neg | Neg |  | 0.00E+00 |
| Culex torrentium | WNV Lineage 1 | 1.00E+07 | Neg | Pos | 22.25 | 1.63E+07 |
| Culex torrentium | WNV Lineage 1 | 1.00E+07 | Neg | Neg |  | 0.00E+00 |
| Culex torrentium | WNV Lineage 1 | 1.00E+07 | Neg | Neg |  | 0.00E+00 |
| Culex torrentium | WNV Lineage 1 | 1.00E+07 | Neg | Neg |  | 0.00E+00 |
| Culex torrentium | WNV Lineage 1 | 1.00E+06 | Neg | Neg |  | 0.00E+00 |
| Culex torrentium | WNV Lineage 1 | 1.00E+06 | Neg | Neg |  | 0.00E+00 |
| Culex torrentium | WNV Lineage 1 | 1.00E+06 | Neg | Neg |  | 0.00E+00 |
| Culex torrentium | WNV Lineage 1 | 1.00E+06 | Neg | Neg |  | 0.00E+00 |
| Culex torrentium | WNV Lineage 1 | 1.00E+06 | Neg | Neg |  | 0.00E+00 |
| Culex torrentium | WNV Lineage 1 | 1.00E+06 | Neg | Neg |  | 0.00E+00 |
| Culex torrentium | WNV Lineage 1 | 1.00E+06 | Neg | Neg |  | 0.00E+00 |
| Culex torrentium | WNV Lineage 1 | 1.00E+06 | Neg | Neg |  | 0.00E+00 |
| Culex torrentium | WNV Lineage 1 | 1.00E+06 | Neg | Neg |  | 0.00E+00 |
| Culex torrentium | WNV Lineage 1 | 1.00E+06 | Neg | Neg |  | 0.00E+00 |
| Culex torrentium | WNV Lineage 1 | 1.00E+06 | Neg | Pos | 36.56 | 1.06E+03 |
| Culex torrentium | WNV Lineage 1 | 1.00E+06 | Neg | Neg |  | 0.00E+00 |
| Culex torrentium | WNV Lineage 1 | 1.00E+06 | Neg | Neg |  | 0.00E+00 |
| Culex torrentium | WNV Lineage 1 | 1.00E+06 | Neg | Neg |  | 0.00E+00 |
| Culex torrentium | WNV Lineage 1 | 1.00E+06 | Neg | Neg |  | 0.00E+00 |
| Culex torrentium | WNV Lineage 1 | 1.00E+05 | Neg | Neg |  | 0.00E+00 |
| Culex torrentium | WNV Lineage 1 | 1.00E+05 | Neg | Neg |  | 0.00E+00 |
| Culex torrentium | WNV Lineage 1 | 1.00E+05 | Neg | Neg |  | 0.00E+00 |
| Culex torrentium | WNV Lineage 1 | 1.00E+05 | Neg | Neg |  | 0.00E+00 |
| Culex torrentium | WNV Lineage 1 | 1.00E+05 | Neg | Neg |  | 0.00E+00 |
| Culex torrentium | WNV Lineage 1 | 1.00E+05 | Neg | Pos | 38.24 | 3.43E+02 |
| Culex torrentium | WNV Lineage 1 | 1.00E+05 | Neg | Neg |  | 0.00E+00 |
| Culex torrentium | WNV Lineage 1 | 1.00E+05 | Neg | Neg |  | 0.00E+00 |
| Culex torrentium | WNV Lineage 1 | 1.00E+05 | Neg | Neg |  | 0.00E+00 |
| Culex torrentium | WNV Lineage 1 | 1.00E+05 | Neg | Pos | 35.64 | 1.98E+03 |
| Culex torrentium | WNV Lineage 1 | 1.00E+05 | Neg | Pos | 36.99 | 7.97E+02 |
| Culex torrentium | WNV Lineage 1 | 1.00E+05 | Neg | Neg |  | 0.00E+00 |
| Culex torrentium | WNV Lineage 1 | 1.00E+05 | Neg | Neg |  | 0.00E+00 |
| Culex torrentium | WNV Lineage 1 | 1.00E+05 | Neg | Neg |  | 0.00E+00 |
| Culex torrentium | WNV Lineage 1 | 1.00E+05 | Neg | Neg |  | 0.00E+00 |
| Culex pipiens biotype pipiens | WNV Lineage 1 | 1.00E+06 | Neg | Neg |  | 0.00E+00 |
| Culex pipiens biotype pipiens | WNV Lineage 1 | 1.00E+06 | Neg | Neg |  | 0.00E+00 |
| Culex pipiens biotype pipiens | WNV Lineage 1 | 1.00E+06 | Neg | Neg |  | 0.00E+00 |
| Culex pipiens biotype pipiens | WNV Lineage 1 | 1.00E+06 | Neg | Pos | 35.91 | 1.65E+03 |
| Culex pipiens biotype pipiens | WNV Lineage 1 | 1.00E+06 | Neg | Neg |  | 0.00E+00 |
| Culex pipiens biotype pipiens | WNV Lineage 1 | 1.00E+06 | Neg | Neg |  | 0.00E+00 |
| Culex pipiens biotype pipiens | WNV Lineage 1 | 1.00E+06 | Neg | Neg |  | 0.00E+00 |
| Culex pipiens biotype pipiens | WNV Lineage 1 | 1.00E+06 | Neg | Neg |  | 0.00E+00 |
| Culex pipiens biotype pipiens | WNV Lineage 1 | 1.00E+06 | Neg | Neg |  | 0.00E+00 |
| Culex pipiens biotype pipiens | WNV Lineage 1 | 1.00E+06 | Neg | Neg |  | 0.00E+00 |
| Culex pipiens biotype pipiens | WNV Lineage 1 | 1.00E+06 | Neg | Pos | 35.22 | 2.62E+03 |
| Culex pipiens biotype pipiens | WNV Lineage 1 | 1.00E+06 | Neg | Neg |  | 0.00E+00 |
| Culex pipiens biotype pipiens | WNV Lineage 1 | 1.00E+06 | Neg | Neg |  | 0.00E+00 |
| Culex pipiens biotype pipiens | WNV Lineage 1 | 1.00E+06 | Neg | Neg |  | 0.00E+00 |
| Culex pipiens biotype pipiens | WNV Lineage 1 | 1.00E+07 | Neg | Neg |  | 0.00E+00 |
| Culex pipiens biotype pipiens | WNV Lineage 1 | 1.00E+07 | Neg | Pos | 33.07 | 1.12E+04 |
| Culex pipiens biotype pipiens | WNV Lineage 1 | 1.00E+07 | Neg | Pos | 35.09 | 2.88E+03 |
| Culex pipiens biotype pipiens | WNV Lineage 1 | 1.00E+07 | Neg | Neg |  | 0.00E+00 |
| Culex pipiens biotype pipiens | WNV Lineage 1 | 1.00E+07 | Neg | Pos | 36.16 | 1.40E+03 |
| Culex pipiens biotype pipiens | WNV Lineage 1 | 1.00E+07 | Neg | Neg |  | 0.00E+00 |
| Culex pipiens biotype pipiens | WNV Lineage 1 | 1.00E+07 | Neg | Neg |  | 0.00E+00 |
| Culex pipiens biotype pipiens | WNV Lineage 1 | 1.00E+07 | Neg | Pos | 37.3 | 6.48E+02 |
| Culex pipiens biotype pipiens | WNV Lineage 1 | 1.00E+07 | Neg | Neg |  | 0.00E+00 |
| Culex pipiens biotype pipiens | WNV Lineage 1 | 1.00E+07 | Neg | Neg |  | 0.00E+00 |
| Culex pipiens biotype pipiens | WNV Lineage 1 | 1.00E+07 | Neg | Neg |  | 0.00E+00 |
| Culex pipiens biotype pipiens | WNV Lineage 1 | 1.00E+07 | Neg | Neg |  | 0.00E+00 |
| Culex pipiens biotype pipiens | WNV Lineage 1 | 1.00E+07 | Neg | Neg |  | 0.00E+00 |
| Culex pipiens biotype pipiens | WNV Lineage 1 | 1.00E+07 | Neg | Neg |  | 0.00E+00 |
| Culex pipiens biotype pipiens | WNV Lineage 1 | 1.00E+07 | Neg | Pos | 36.69 | 9.77E+02 |
| Culex pipiens biotype pipiens | WNV Lineage 1 | 1.00E+07 | Neg | Neg |  | 0.00E+00 |
| Culex pipiens biotype pipiens | WNV Lineage 1 | 1.00E+07 | Neg | Neg |  | 0.00E+00 |
| Culex pipiens biotype pipiens | WNV Lineage 1 | 1.00E+07 | Neg | Neg |  | 0.00E+00 |
| Culex pipiens biotype pipiens | WNV Lineage 1 | 1.00E+05 | Neg | Neg |  | 0.00E+00 |
| Culex pipiens biotype pipiens | WNV Lineage 1 | 1.00E+05 | Neg | Neg |  | 0.00E+00 |
| Culex pipiens biotype pipiens | WNV Lineage 1 | 1.00E+05 | Neg | Neg |  | 0.00E+00 |
| Culex pipiens biotype pipiens | WNV Lineage 1 | 1.00E+05 | Neg | Neg |  | 0.00E+00 |
| Culex pipiens biotype pipiens | WNV Lineage 1 | 1.00E+05 | Neg | Neg |  | 0.00E+00 |
| Culex pipiens biotype pipiens | WNV Lineage 1 | 1.00E+05 | Neg | Neg |  | 0.00E+00 |
| Culex pipiens biotype pipiens | WNV Lineage 1 | 1.00E+05 | Neg | Neg |  | 0.00E+00 |
| Culex pipiens biotype pipiens | WNV Lineage 1 | 1.00E+05 | Neg | Neg |  | 0.00E+00 |
| Culex pipiens biotype pipiens | WNV Lineage 1 | 1.00E+05 | Neg | Neg |  | 0.00E+00 |
| Culex pipiens biotype pipiens | WNV Lineage 1 | 1.00E+05 | Neg | Pos | 35.56 | 2.08E+03 |
| Culex pipiens biotype pipiens | WNV Lineage 1 | 1.00E+05 | Neg | Neg |  | 0.00E+00 |
| Culex pipiens biotype pipiens | WNV Lineage 1 | 1.00E+05 | Neg | Neg |  | 0.00E+00 |
| Culex pipiens biotype pipiens | WNV Lineage 1 | 1.00E+05 | Neg | Neg |  | 0.00E+00 |
| Culex pipiens biotype pipiens | WNV Lineage 1 | 1.00E+05 | Neg | Neg |  | 0.00E+00 |
| Culex pipiens biotype pipiens | WNV Lineage 1 | 1.00E+05 | Neg | Neg |  | 0.00E+00 |
| Culex pipiens biotype pipiens | WNV Lineage 1 | 1.00E+06 | Neg | Pos | 38.11 | 3.75E+02 |
| Culex pipiens biotype pipiens | WNV Lineage 1 | 1.00E+06 | Neg | Neg |  | 0.00E+00 |
| Culex pipiens biotype pipiens | WNV Lineage 1 | 1.00E+06 | Neg | Neg |  | 0.00E+00 |
| Culex pipiens biotype pipiens | WNV Lineage 1 | 1.00E+06 | Neg | Neg |  | 0.00E+00 |
| Culex pipiens biotype pipiens | WNV Lineage 1 | 1.00E+06 | Neg | Neg |  | 0.00E+00 |
| Culex pipiens biotype pipiens | WNV Lineage 1 | 1.00E+06 | Neg | Neg |  | 0.00E+00 |
| Culex pipiens biotype pipiens | WNV Lineage 1 | 1.00E+06 | Neg | Neg |  | 0.00E+00 |
| Culex pipiens biotype pipiens | WNV Lineage 1 | 1.00E+06 | Neg | Neg |  | 0.00E+00 |
| Culex pipiens biotype pipiens | WNV Lineage 1 | 1.00E+06 | Neg | Neg |  | 0.00E+00 |
| Culex pipiens biotype pipiens | WNV Lineage 1 | 1.00E+06 | Neg | Neg |  | 0.00E+00 |
| Culex pipiens biotype pipiens | WNV Lineage 1 | 1.00E+06 | Neg | Neg |  | 0.00E+00 |
| Culex pipiens biotype pipiens | WNV Lineage 1 | 1.00E+06 | Neg | Neg |  | 0.00E+00 |
| Culex pipiens biotype pipiens | WNV Lineage 1 | 1.00E+06 | Neg | Neg |  | 0.00E+00 |
| Culex pipiens biotype pipiens | WNV Lineage 1 | 1.00E+06 | Neg | Pos | 34.49 | 2.63E+03 |
| Culex pipiens biotype pipiens | WNV Lineage 1 | 1.00E+06 | Neg | Pos | 37.71 | 2.26E+02 |
| Culex pipiens biotype pipiens | WNV Lineage 1 | 1.00E+06 | Neg | Neg |  | 0.00E+00 |
| Culex pipiens biotype pipiens | WNV Lineage 1 | 1.00E+05 | Neg | Neg |  | 0.00E+00 |
| Culex pipiens biotype pipiens | WNV Lineage 1 | 1.00E+05 | Neg | Neg |  | 0.00E+00 |
| Culex pipiens biotype pipiens | WNV Lineage 1 | 1.00E+05 | Neg | Neg |  | 0.00E+00 |
| Culex pipiens biotype pipiens | WNV Lineage 1 | 1.00E+05 | Neg | Neg |  | 0.00E+00 |
| Culex pipiens biotype pipiens | WNV Lineage 1 | 1.00E+05 | Neg | Neg |  | 0.00E+00 |
| Culex pipiens biotype pipiens | WNV Lineage 1 | 1.00E+05 | Neg | Pos | 35.08 | 1.68E+03 |
| Culex pipiens biotype pipiens | WNV Lineage 1 | 1.00E+05 | Neg | Neg |  | 0.00E+00 |
| Culex pipiens biotype pipiens | WNV Lineage 1 | 1.00E+05 | Neg | Neg |  | 0.00E+00 |
| Culex pipiens biotype pipiens | WNV Lineage 1 | 1.00E+05 | Neg | Neg |  | 0.00E+00 |
| Culex pipiens biotype pipiens | WNV Lineage 1 | 1.00E+05 | Neg | Neg |  | 0.00E+00 |
| Culex pipiens biotype pipiens | WNV Lineage 1 | 1.00E+05 | Neg | Neg |  | 0.00E+00 |
| Culex pipiens biotype pipiens | WNV Lineage 1 | 1.00E+05 | Neg | Pos | 36.57 | 5.36E+02 |
| Culex pipiens biotype pipiens | WNV Lineage 1 | 1.00E+05 | Neg | Neg |  | 0.00E+00 |
| Culex pipiens biotype pipiens | WNV Lineage 1 | 1.00E+05 | Neg | Neg |  | 0.00E+00 |
| Culex pipiens biotype pipiens | WNV Lineage 1 | 1.00E+05 | Neg | Neg |  | 0.00E+00 |
| Culex torrentium | WNV Lineage 2 | 1.00E+05 | Neg | Neg |  | 0.00E+00 |
| Culex torrentium | WNV Lineage 2 | 1.00E+05 | Neg | Neg |  | 0.00E+00 |
| Culex torrentium | WNV Lineage 2 | 1.00E+05 | Neg | Neg |  | 0.00E+00 |
| Aedes albopictus | WNV Lineage 1 | 1.00E+06 | Neg | Neg |  | 0.00E+00 |
| Aedes albopictus | WNV Lineage 1 | 1.00E+06 | Neg | Neg |  | 0.00E+00 |
| Aedes albopictus | WNV Lineage 1 | 1.00E+06 | Neg | Neg |  | 0.00E+00 |
| Aedes albopictus | WNV Lineage 1 | 1.00E+06 | Neg | Neg |  | 0.00E+00 |
| Aedes albopictus | WNV Lineage 1 | 1.00E+06 | Neg | Pos | 37.24 | 3.22E+02 |
| Aedes albopictus | WNV Lineage 1 | 1.00E+06 | Neg | Neg |  | 0.00E+00 |
| Aedes albopictus | WNV Lineage 1 | 1.00E+06 | Neg | Neg |  | 0.00E+00 |
| Aedes albopictus | WNV Lineage 1 | 1.00E+06 | Neg | Neg |  | 0.00E+00 |
| Aedes albopictus | WNV Lineage 1 | 1.00E+06 | Neg | Pos | 36.88 | 4.26E+02 |
| Culex pipiens biotype pipiens | WNV Lineage 2 | 1.00E+05 | Neg | Pos | 36.44 | 5.94E+02 |
| Culex pipiens biotype pipiens | WNV Lineage 2 | 1.00E+05 | Neg | Neg |  | 0.00E+00 |
| Culex pipiens biotype pipiens | WNV Lineage 2 | 1.00E+05 | Neg | Pos | 36.4 | 6.12E+02 |
| Culex pipiens biotype pipiens | WNV Lineage 2 | 1.00E+05 | Neg | Neg |  | 0.00E+00 |
| Culex pipiens biotype pipiens | WNV Lineage 2 | 1.00E+05 | Neg | Neg |  | 0.00E+00 |
| Culex pipiens biotype pipiens | WNV Lineage 2 | 1.00E+05 | Neg | Neg |  | 0.00E+00 |
| Culex pipiens biotype pipiens | WNV Lineage 2 | 1.00E+05 | Neg | Neg |  | 0.00E+00 |
| Culex pipiens biotype pipiens | WNV Lineage 2 | 1.00E+05 | Neg | Neg |  | 0.00E+00 |
| Culex pipiens biotype pipiens | WNV Lineage 2 | 1.00E+05 | Neg | Neg |  | 0.00E+00 |
| Culex pipiens biotype pipiens | WNV Lineage 2 | 1.00E+05 | Neg | Neg |  | 0.00E+00 |
| Culex pipiens biotype pipiens | WNV Lineage 2 | 1.00E+05 | Neg | Neg |  | 0.00E+00 |
| Culex pipiens biotype pipiens | WNV Lineage 2 | 1.00E+05 | Neg | Neg |  | 0.00E+00 |
| Culex pipiens biotype pipiens | WNV Lineage 2 | 1.00E+05 | Neg | Pos | 36.75 | 4.68E+02 |
| Culex pipiens biotype pipiens | WNV Lineage 2 | 1.00E+05 | Neg | Neg |  | 0.00E+00 |
| Culex pipiens biotype pipiens | WNV Lineage 2 | 1.00E+05 | Neg | Neg |  | 0.00E+00 |
| Culex pipiens biotype pipiens | WNV Lineage 2 | 1.00E+05 | Neg | Neg |  | 0.00E+00 |
| Culex pipiens biotype pipiens | WNV Lineage 2 | 1.00E+05 | Neg | Neg |  | 0.00E+00 |
| Culex pipiens biotype pipiens | WNV Lineage 2 | 1.00E+05 | Neg | Neg |  | 0.00E+00 |
| Culex pipiens biotype pipiens | WNV Lineage 2 | 1.00E+05 | Neg | Neg |  | 0.00E+00 |
| Culex pipiens biotype pipiens | WNV Lineage 2 | 1.00E+05 | Neg | Neg |  | 0.00E+00 |
| Culex pipiens biotype pipiens | WNV Lineage 2 | 1.00E+05 | Neg | Neg |  | 0.00E+00 |
| Culex pipiens biotype pipiens | WNV Lineage 2 | 1.00E+05 | Neg | Neg |  | 0.00E+00 |
| Culex pipiens biotype pipiens | WNV Lineage 2 | 1.00E+05 | Neg | Pos | 19.27 | 2.87E+08 |
| Culex pipiens biotype pipiens | WNV Lineage 2 | 1.00E+05 | Neg | Neg |  | 0.00E+00 |
| Culex pipiens biotype pipiens | WNV Lineage 2 | 1.00E+05 | Pos | Pos | 18.53 | 5.04E+08 |
| Culex pipiens biotype pipiens | WNV Lineage 2 | 1.00E+05 | Neg | Pos | 21.58 | 4.91E+07 |
| Culex pipiens biotype pipiens | WNV Lineage 2 | 1.00E+05 | Neg | Pos | 35.99 | 8.42E+02 |
| Culex pipiens biotype pipiens | WNV Lineage 2 | 1.00E+05 | Neg | Neg |  | 0.00E+00 |
| Culex pipiens biotype pipiens | WNV Lineage 2 | 1.00E+05 | Neg | Neg |  | 0.00E+00 |
